# A Global Thermodynamic-Kinetic Model Capturing the Hallmarks of Liquid-Liquid Phase Separation and Amyloid Aggregation

**DOI:** 10.1101/2025.05.23.655849

**Authors:** Kamal Bhandari, Yunxiang Sun, Huayuan Tang, Pu Chu Ke, Feng Ding

## Abstract

Aberrant aggregation of proteins into amyloid fibrils is associated with numerous neurodegenerative, systemic and metabolic diseases. Amyloidogenic proteins undergo spontaneous liquid-liquid phase separation (LLPS), rapidly forming protein-rich condensates prior to fibrillization. However, the exact effects of LLPS on amyloid aggregation remain unclear as contrasting fibrillization-promotion, inhibition and even biphasic effects have been reported in the literature. In this study, we integrate LLPS-induced heterogeneity of protein concentrations into a thermodynamic-kinetic model of amyloid aggregation. We adopt the phase transition theory and introduce protein condensates as an additional protein state alongside non-interacting monomers, oligomers and fibrils. Oligomerization and fibrillization can occur both in the protein-rich condensates and the protein-poor solution. This model allows us to derive the time evolution of different states – monomers, condensates, oligomers, and fibrils – spanning a wide range of concentrations, and determine how model parameters related to LLPS, fibrillization, and oligomerization influence fibrillization kinetics. Using this global model, we resolve the seemingly contradictory effects of LLPS on fibrillization. We expect the developed thermodynamic-kinetic model of LLPS, and amyloid aggregation will help advance our understanding, modulation, and mitigation of pathological aggregation processes in amyloid diseases.

**TOC:** 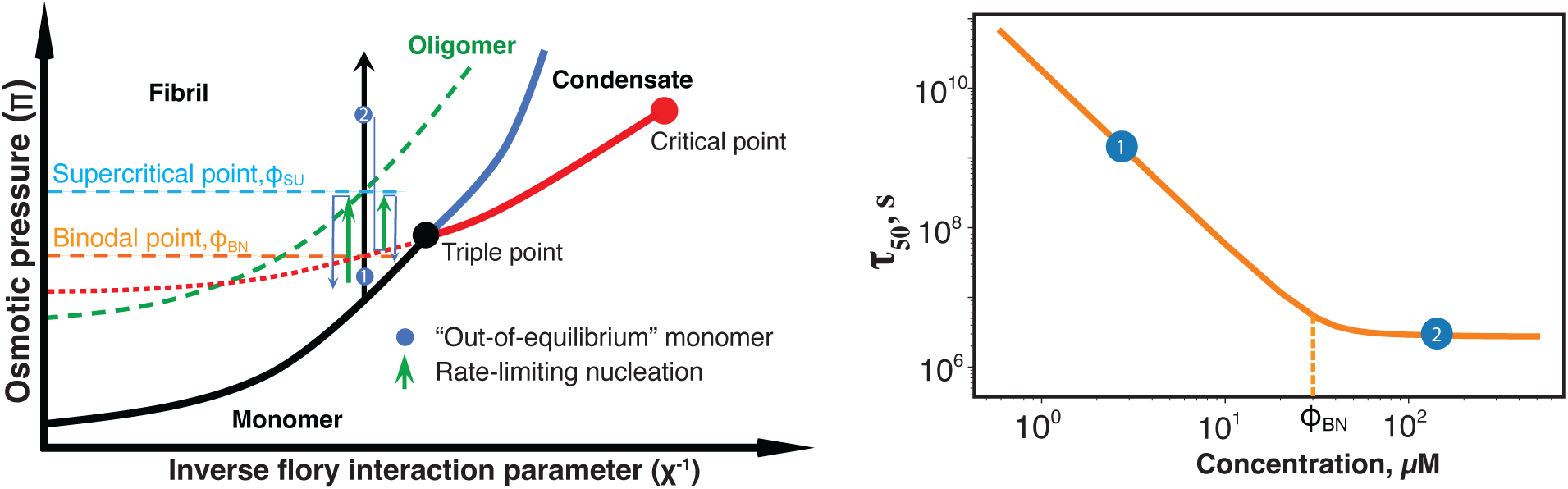

## Introduction

A common pathological hallmark of neurodegenerative disorders such as Alzheimer’s disease (AD), Parkinson’s disease (PD), amyotrophic lateral sclerosis (ALS) as well as metabolic type-2 diabetes (T2D) is the deposition of aggregated proteins in the form of amyloid fibrils in tissues and cells (1–5). These diseases pose significant challenges to both patients and healthcare systems globally due to a lack of understanding of their pathophysiology and hence a lack of mitigation strategies. With recent successes of antibody therapeutics in delaying the progression of early-stage AD by targeting and clearing amyloid aggregates (6), there is a renewed interest in the “amyloid hypothesis”, which suggests that amyloid aggregation is a major cause of neurodegeneration and eventual cognitive and motor deficits. Thus, a better understanding of the mechanisms of amyloid aggregation is crucial for the development of effective approaches to modulate and mitigate amyloid aggregation and their entailed neurotoxicity.

Recent technical advancement in cryo-electron microscopy and solid-state nuclear magnetic resonance has enabled high-resolution imaging of the fibrillar structures of many amyloid proteins (7–10), revealing a common cross-β structure where monomeric proteins stack into parallel, in-register intermolecular β-sheets with the β-strands aligned perpendicularly to the fibril axis and packed against each other to render the cross-β core (11–16). Interestingly, fibrils formed by the same amyloid proteins may display distinct morphologies due to different backbone arrangements of composite proteins in the protofilaments and/or packing between protofilaments (8, 17, 18). Despite the diversity in sequence and structure, the aggregation processes from monomers to fibrils share similar sigmoidal kinetics for all amyloid proteins (11), characterized by an initial lag phase followed by a rapid exponential-like growth phase and a final plateau phase due to saturation and depletion of monomers. Soluble aggregation intermediates, rather than mature fibrils, are increasingly recognized as key initiators in eliciting cytotoxicity in amyloid diseases (19–21).

Recent studies have revealed an intricate relationship between liquid-liquid phase separation (LLPS) and amyloid aggregation. LLPS denotes the spontaneous separation of a homogeneous solution into two phases: a dense phase enriched with biomolecules and a solution phase depleted of them, exhibiting contrasting physical properties such as viscosity and diffusivity. Examples of the physiological or pathological condensates include nucleoli, promyelocytic leukemia nuclear bodies, and stress granules (22–25). These condensates are liquid-like, with composite biomolecules forming a network facilitated by weak and dynamic intermolecular interactions (26–30). Increasing experimental studies have revealed that amyloid proteins, including amyloid-β (Aβ) (31), α-synuclein (32), tau (33, 34), islet amyloid polypeptide (IAPP) (35, 36), insulin A-chain fragment (37), and TDP-43 (38), undergo LLPS to render protein condensates. The condensates typically assume the morphology of spheres and are often referred to as “droplets” in the literature, providing temporal and spatial organizations of amyloid proteins *in vivo*. Using coarse-grained molecular dynamics simulations (36), Xing et al. demonstrated both LLPS and fibrillization *in silico*, providing a unified picture of amyloid aggregation for a wide range of concentrations within the framework of LLPS. LLPS-driven condensates not only serve as dynamic compartments within the cell but also act as potent modulators of amyloid aggregation kinetics through concentration partitioning. It has been postulated that the elevated concentration of amyloidogenic proteins within these condensates promotes nucleation (39–41), thereby facilitating the formation of toxic oligomers and amyloid fibrils. On the other hand, experimental studies have also shown that the formation of condensates significantly hinder the nucleation and growth of amyloid fibrils (42–44). Moreover, a concentration-dependent biphasic effect on α-synuclein fibrillization by a droplet-promoting salt was reported (45), where the initial fibrillization-promotion effect at low to moderate salt concentrations was reversed above a certain threshold. Hence, the effect of protein condensate formation via LLPS on amyloid aggregation is likely nonlinear. Understanding both LLPS and amyloid aggregation as well as their interplay marks a key research frontier of amyloid aggregation with implications for therapeutic design and intervention against amyloid diseases.

Nucleation-dependent polymerization models are commonly used to describe amyloid aggregation kinetics, where the initial lag phase originates from the formation of critical nuclei via the self-assembly of monomers in the solution, known as primary nucleation (46). Variations of the classical nucleation theory, including the surface-catalyzed secondary nucleation and/or fragmentation (15, 47), supercritical concentration due to the monomer-oligomer equilibrium (48), and the nucleated conformational conversion (49) have been developed to capture the concentration-dependence of amyloid aggregation, such as the saturation of aggregation kinetics at high concentrations (15, 49–53). However, these nucleation-dependent models do not account for LLPS-induced concentration heterogeneity, and instead treat protein solutions as homogeneous away from reality. The oligomer intermediates, either explicitly or implicitly considered in these models, are size-limited up to tens of proteins with nano-meters in dimension and, thus, do not correspond to protein droplets which can be as large as micro-meters as observed experimentally (31–35). There is a *critical need* to develop an aggregation model that integrates LLPS and fibrillization to accurately capture the whole aggregation kinetics. By drawing analogies between LLPS and gas-liquid phase transition as well as between highly ordered amyloid fibrils and solid crystals (54), we previously proposed a phase diagram for non-interacting monomers, droplets and fibrils and postulated that amyloid aggregation could be regarded as the equilibration from an initial out-of-equilibrium state of monomer solution to the equilibrium state of stable fibrils coexisting with monomers and/or droplets. Here, we adopted the proposed phase transition theory and developed an analytical model for amyloid aggregation that captured LLPS while incorporating LLPS-induced heterogeneity of protein solutions.

Specifically, we used the classical nucleation theory to model the formation of both droplets and fibrils and assumed that LLPS is orders-of-magnitude faster than fibrillization. In our model, oligomers were regarded as fibrillization intermediates. Both oligomers and fibril seeds could form within the protein-rich condensates and protein-poor solution. The droplet-depletion induced exposure of preformed fibrils in the droplets was modeled by a mean-field approach. This model allowed us to examine amyloid aggregation kinetics over a wide range of protein concentrations, crossing the binodal concentration of LLPS where the spontaneous formation of protein droplets occurs. In our model, fibrillization accelerated and the aggregation lag-time or half-time decreased following a power-law scaling with protein concentration below the binodal point. But above the binodal point, the trend rapidly reached a plateau as observed experimentally (36). Importantly, by differentiating experimentally-used droplet-promoting approaches into those primarily impacting proteins in either the solution, the droplets, or both, we were able to rationalize the seemingly contradicting effects of droplet formation on fibrillization as reported in the literature (55). We also analyzed the impact of fragmentation and secondary nucleation in the aggregation kinetics of our model. With tunable parameters capturing different protein-solution systems, our model is expected to be applicable to delineating amyloid aggregation behaviors under various conditions and identifying efficient approaches to modulate and mitigate pathological aggregation processes in combating amyloid diseases.

## Results and Discussion

### Modeling both LLPS and fibrillization with a global nucleation-dependent theory

LLPS is a rapid process where biomolecules above a threshold concentration quickly form initial droplets within seconds to minutes (25). In contrast, the fibrillization of amyloid proteins is a much slower process, which occurs over hours to days *in vitro* at higher concentrations (56) and takes decades to develop and manifest clinically. These differences in timescales can be attributed to the different free-energy barriers associated with the two phase-transition processes (54). Specifically, the fluid-like, highly dynamic droplets of LLPS are stabilized by weak non-specific intermolecular interactions, and proteins are not required to undergo major conformational changes upon forming droplets. On the other hand, nucleation of amyloid proteins involves major conformational changes from either intrinsically disordered or otherwise folded states to a fibrillar structure stabilized by extensive intermolecular hydrogen bonds, leading to a significant conformational entropy loss.

With the clear separation of timescales between LLPS and fibrillization, we expect the de-mixing of proteins to occur first in solution, reaching the monomer-droplet equilibrium with fast exchanges of monomers between the protein-poor solution and protein-rich droplets (step i, **Fig. 1A**). As observed experimentally, fibril seeds can nucleate out of both protein-poor and protein-rich phases (steps ii & iii, **Fig. 1A**). Starting from the nucleated seeds, fibrils rapidly grew via binding and seeded conformational conversion of protein monomers available in both phases. The removal of proteins from both phases into fibrils caused a depletion of droplets because of the rapidly shifting monomer-droplet equilibrium, resulting in the exposure of fibrils initially nucleated in the droplets to the protein-poor solution (step iv, **Fig. 1A**). With the complete depletion of droplets, fibrils grew only from the homogenous phase of monomers until an equilibrium between the monomers and fibrils was reached. Next, we discuss how to model each of these steps.

**Figure 1.**
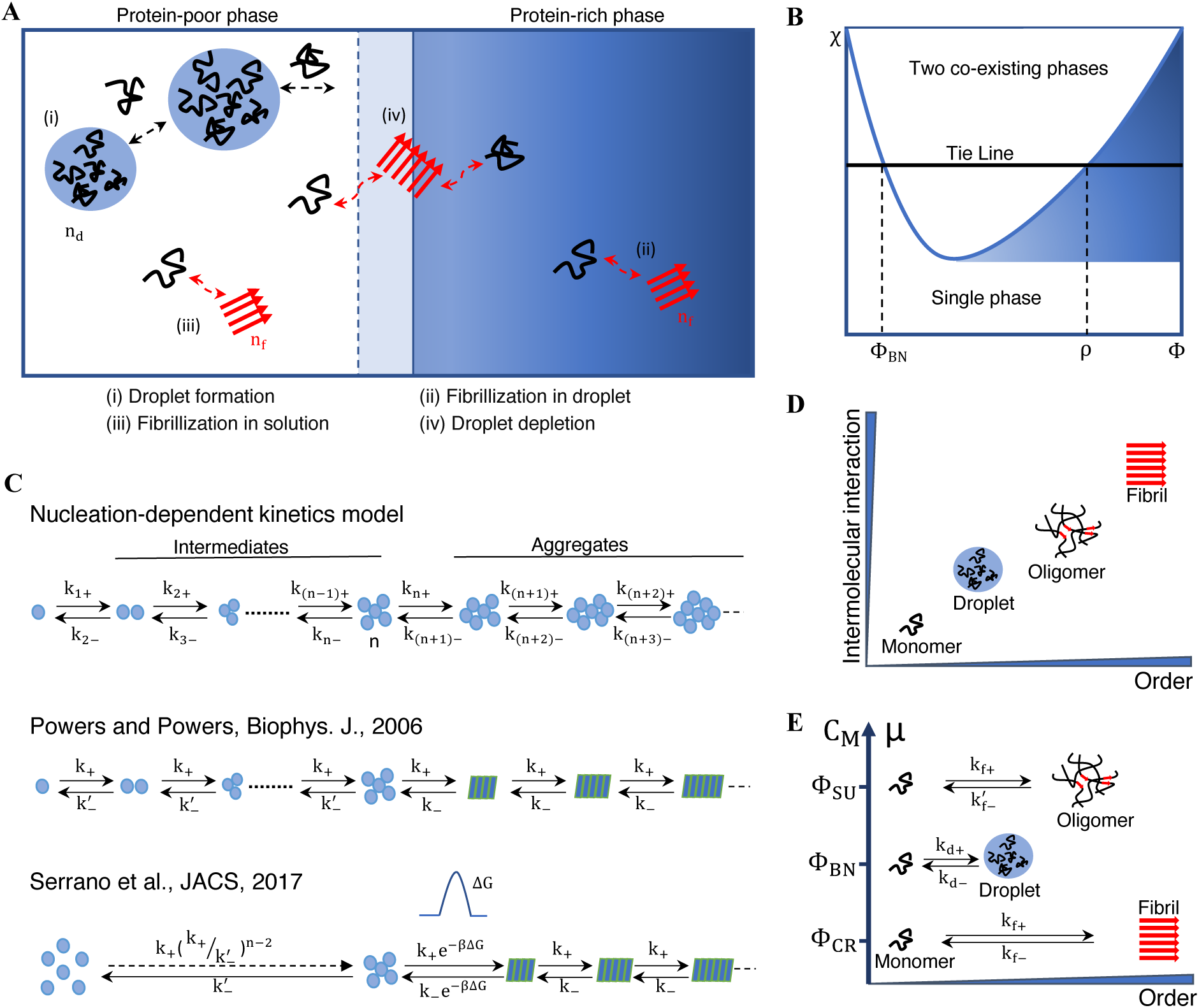
Modeling both LLPS and fibrillization in amyloid protein aggregation. (**A**) Schematic diagram of four processes considered in our model, including droplet formation (i), fibrillization in droplets (ii) and in solution (iii), and droplet-depletion induced exposure of pre-formed fibrils in the droplets (iv). Here, n_d_ and n_f_ represent critical nucleus of droplets and fibrils, correspondingly. Black curly lines represent monomers, and solid red arrows represent oligomers or fibrils. Dashed black arrows represent the monomer exchange between the solution and droplet during droplets growth. Dashed red arrows represent the monomer accumulation or addition to nucleate the oligomer or fibrils. The light blue shaded zone with a dashed boundary represents oligomers or fibrils that are nucleated originally within the droplets but now are exposed to the solution. (**B**) A phase diagram of LLPS shown as a function of the Flory parameter, χ, and concentration, Φ. A tie line with a constant χ denotes a solute-solvent system under a given temperature, and it intercepts the co-existing line at the binodal points, Φ_BN_ < ρ. With Φ in between the two points, an initially homogenous system spontaneously separates into protein-rich droplets at concentration ρ and and a protein-poor solution at the concentration Φ_BN_. Shaded blue region represents the droplet phase as in panel A. (**C**) Typical nucleation-dependent aggregation models depicting the assembly of monomers into various intermediates up to the critical nuclei with size n, and the subsequent growth of aggregates via monomer addition. Powers and Powers (48) differentiated the dissociation rates of intermediates and fibrils (highlighted as rectangle shapes). In Serrano et al.(49), a free-energy barrier was introduced between the nuclei and fibril seeds, associated with the nucleated conformational conversion of oligomers (52). (**D**) A cartoon representation of thermodynamic profiles of four protein states – monomers, droplets, oligomers, and fibrils – in LLPS and fibrillization in terms of intermolecular interaction and structural order. (**E**) Chemical potential μ of proteins in each of the four states in panel **D**. The μ of non-interacting protein monomer solution depends on the concentration C_M_ and the equilibrium concentration of monomers with respect to fibrils (the critical concentration, Φ_CR_), droplets (the binodal concentration, Φ_BN_), and oligomers (the supercritical concentration, Φ_SU_).

LLPS is thermodynamically described by the Flory-Huggins theory (57, 58) (e.g., the schematic phase diagram in **Fig. 1B**). Using a mean-field approximation, the free-energy change associated with mixing polymers and solvents depends on a dimensionless interaction parameter, χ, and the total polymer concentration, Φ. Here, the Flory parameter χ represents the average energy gain of two polymeric units (i.e., the composite amino acids of proteins) excluding their solvent to form close contacts, measured in units of thermal energy KT, where K is the Boltzmann constant, and T is temperature. Based on the second law of thermodynamics, phase separation can occur when polymers exhibit attractive interactions with each other under given solution condition (e.g., temperature, pH, salt concentration, co-solutes, etc.), providing that the corresponding χ exceeds the critical value of approximately 0.5. This threshold is generally met for amyloid proteins capable of self-assembling into fibrils. For an amyloid protein under a given solution condition, its phase separation behavior is determined by the total protein concentration along the tie line with the fixed χ-value (**Fig. 1B**). Spontaneous de-mixing of the solution into protein-rich and protein-poor phases occurs only when the total protein concentration Φ increases above the binodal point, Φ_BN_ (57, 58). In equilibrium, droplets with a high protein concentration of ρ co-exist with the protein-poor solution at the concentration of Φ_BN_.

Similarly to the modeling of fibrillization, we also adopted the classical nucleation theory to model protein droplet formation – where all aggregation species grow or shrink through monomer association and dissociation, and the aggregates of droplets/fibrils are considered stable only after the initial formation of a critical nucleus with the size of n proteins (**Fig. 1C**) (59, 60). Although this model might oversimplify the kinetics of phase separation, especially at high concentrations above the so-called spinodal point where nucleation-independent de-mixing (i.e., the spinodal decomposition) may occur, it can accurately capture the equilibrium partition of amyloid proteins between two co-exiting phases. Since the rates of amyloid aggregation are orders-of-magnitude slower than droplet formation, the equilibrium of LLPS is effectively maintained from the perspective of fibrillization, which is the focus of this study.

In nucleation-dependent aggregation/polymerization models, the aggregation kinetics can be determined by solving the rate equations with given rate constants and initial conditions (59, 60). For instance, by assuming the same association (k_+_) and dissociation (k_–_) rates for all aggregates and a rapid pre-equilibrium of monomers with weakly populated intermediates, a closed-form solution of aggregation kinetics can be obtained (48) and the aggregation half-time t_50_ follows a power-law scaling with the total protein concentration, 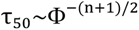. The ratio k_–_/k_+_ defines the critical fibrillization concentration of proteins, Φ_CR_, only above which fibrils are stable. However, this scaling of aggregation kinetics is valid only at low concentrations when the monomers are less stable than intermediates. To capture deviations from the predicted power-law scaling at high concentrations, Powers and Powers (48) explicitly considered the intermediates with a faster dissociation rate, 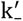 (**Fig. 1C**), and defined a “supercritical concentration”, Φ_SU_ = k’_–_/k_+_, above which protein monomers initially reach a quasi-equilibrium with rapidly formed intermediates before the conversion and growth of fibrils. As a result, the Powers-Powers model captured the slowing down of amyloid aggregation kinetics at high protein concentrations above Φ_SU_. To model the nucleated conformational conversion (52, 61) of oligomers into highly ordered fibril seeds, Serrano et al.(49) introduced a free-energy barrier (ΔG) separating the nuclei and fibril seeds, corresponding to the associated conformational transition (**Fig. 1C**). For simplicity, Serrano et al. adopted a pre-equilibrium approximation for smaller intermediates and assumed that multiple monomers simultaneously assembled into the largest intermediate or critical nucleus, with a corresponding nucleation rate, 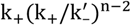. This approximation resulted in a similar slowing down of aggregation kinetics above the supercritical concentration (48).

In this study, we used a similar approximation as in Serrano et al. (49) to model both LLPS and fibrillization. For LLPS, the ratio of droplet dissociation and association rates, k_d–_/k_d+_, denotes the binodal concentration, Φ_BN_, while the ratio k’_d–_/k_d+_ relates to the formation of droplet intermediates. Given the fast de-mixing of LLPS and minimal conformational changes of amyloid proteins upon forming droplets, we set the corresponding ΔG_LLPS_ zero and counted the LLPS nuclei with size n_d_as part of the droplets. In contrast, the fibrillization nuclei with size n_f_represents amyloid oligomers, which have less ordered conformations than fibrils but with significantly stronger intermolecular interactions than droplets (**Fig. 1D**). Therefore, we considered these amyloid oligomers distinct from fibrils with a non-zero free-energy barrier ΔG, and the corresponding supercritical concentration is defined as Φ_SU_ = k’_f–_/k_f+_ and critical fibrillization concentration Φ_CR_ = k_f–_/k_f+_.

Here, Φ_CR_, Φ_SU_, and Φ_BN_ correspond to the protein monomer concentrations in equilibrium with fibrils, oligomers, and droplets, respectively. The chemical potential of non-interacting protein monomers in solution is proportional to the natural logarithm of their concentration (Φ), μ = μ_0_ + kT ln(Φ) – i.e., the lower the concentration, the lower the chemical potential. Since two states at diffusive equilibrium have the same chemical potential, the chemical potential of composite proteins in fibrils, oligomers, or droplets is thus proportional to the logarithm of Φ_CR_, Φ_SU_, and Φ_BN_, correspondingly (**Fig. 1E**). The highly ordered fibril state is the most thermally stable with the lowest chemical potential. This strong stability arises from extensive hydrogen bonds formed between neighboring proteins in the fibrils, even though the entropy in this state is the lowest among the three protein-interacting states. Compared to droplets, proteins in oligomers (i.e., the fibrillization intermediates and nuclei) possess a lower enthalpy with stronger inter-protein interactions, but also a lower entropy due to their greater structural order and less conformational flexibility. Therefore, the relative chemical potentials of oligomers and droplets depend on whether enthalpic gains or entropic penalties dominate under a given condition, including the solution environment and specific proteins under study.

We distinguished oligomers and fibrils formed in the high-density liquid phase of protein droplets from those formed out of the low-density liquid phase of non-interacting protein monomer solution. For simplicity, we treated droplets of different sizes as a single phase for fibrillization. Due to rapid monomer-droplet equilibration, the formation of oligomers and fibrils inside the droplets does not reduce the high protein concentration but rather decreases the total droplet mass or volume. With shrinking total droplet volume, the oligomers and fibrils initially formed in the droplets became partially exposed to the monomer solution, and they grew by the addition of proteins in both protein-rich and poor phases (step iv, **Fig. 1A**). To model this effect, we assumed that the droplet-depletion induced exposure was equally applied to all oligomers and fibrils initially formed in the droplets. Taken together, our model can be described by the following simplified rate equations, with detailed derivations and descriptions provided in the Supplementary Information (**SI**):

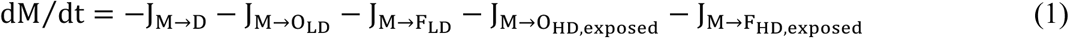

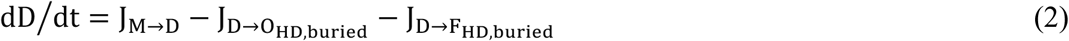

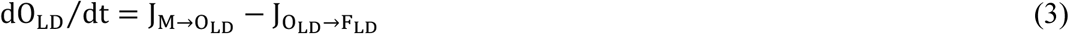

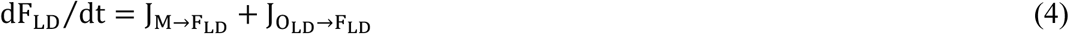

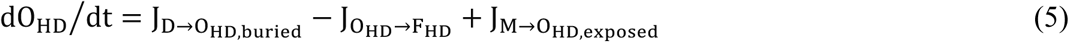

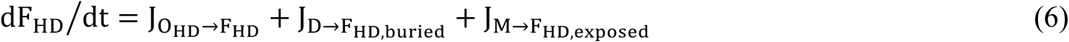

where parameters M and D represent the concentrations of total proteins in the non-interacting monomer solution and protein-rich droplets, respectively; while O and F stand for the total protein concentration of oligomers and fibrils with the subscripts LD and HD denoting their origins from low-density and high-density liquid phases of the heterogeneous protein solution, respectively. J represents the net flux of proteins between different states, as indicated by arrows in the subscript. The notations of *buried* and *exposed* were added for oligomers and fibrils initially formed inside the droplets to indicate their droplet-depletion induced exposure. We used the volume fraction of total droplets D/D_max_ to model the average exposure, where D_max_ denotes the maximum protein concentration of droplets reached during the initial monomer-droplet pre-equilibration.

Fibril growth inside droplets thermodynamically depends on the chemical potential difference of proteins in fibrils and in droplets. The chemical potential of fibrils mostly depends on inter-protein interactions in the cross-β structures, and likely, does not differ significantly in the protein-rich or poor phases. The chemical potential of droplets, on the other hand, equals to that of non-interacting monomer solution at the concentration of Φ_BN_. Therefore, fibril growth in droplets with a high concentration of ρ is thermodynamically equivalent to the growth in solution with a protein concentration of Φ_BN_, but not necessarily kinetically. To account for this effect, the fibril growth and shrinkage rates inside the droplets are proportional to those in solution, by a proportionality coefficient σ, which quantifies the enhancement (σ>1) or suppression (σ<1) of fibrillization rates due to phase separation by capturing the differences in kinetic parameters between the high-density and low-density phases. The fibril growth rate equals to the association rate times the monomer concentration, and the fibril shrinkage rate is the same as the dissociation rate. Hence, ρk_HD, f+_ = σΦ_BN_k_LD, f+_ and k_HD, f–_ = σk_LD, f–_. The same relationship also applies to the formation of oligomers with k’_HD, f–_ = σk’_LD, f–_. Similarly, the free-energy barriers for fibril nucleation in the droplets (ΔG_HD_) and in solution (ΔG_LD_) can be different, and their difference ΔΔG = ΔG_HD_ - ΔG_LD_ determines the corresponding differential effects in the two phases of LLPS. Using this global model of LLPS and amyloid aggregation (Eqs. 1–6), we next investigated the effects of LLPS on oligomerization and fibrillization.

### Concentration dependence of the global LLPS-fibrillization model

The rate-mass equations Eqs. 1–6 include multiple free parameters that can be tailored for different protein-solvent systems to study the corresponding aggregation kinetics. We started with a simple model system with all the parameter values listed in **Table 1** and a detailed justification for these values given in the **SI**. In this case, we assumed the same fibrillization kinetics in the droplets as in solution (σ=1 and ΔΔG=0) that oligomers were less stable than droplets (Φ_BN_ < Φ_SU_). Given the assigned parameters, we integrated the differential equations of Eqs. 1–6 and computed the time evolutions of different species - monomer, droplets, oligomers, and fibrils. We performed calculations for a broad range of total protein concentrations to investigate the concentration dependence of both LLPS and fibrillization (**Fig. 2**).

**Figure 2.**
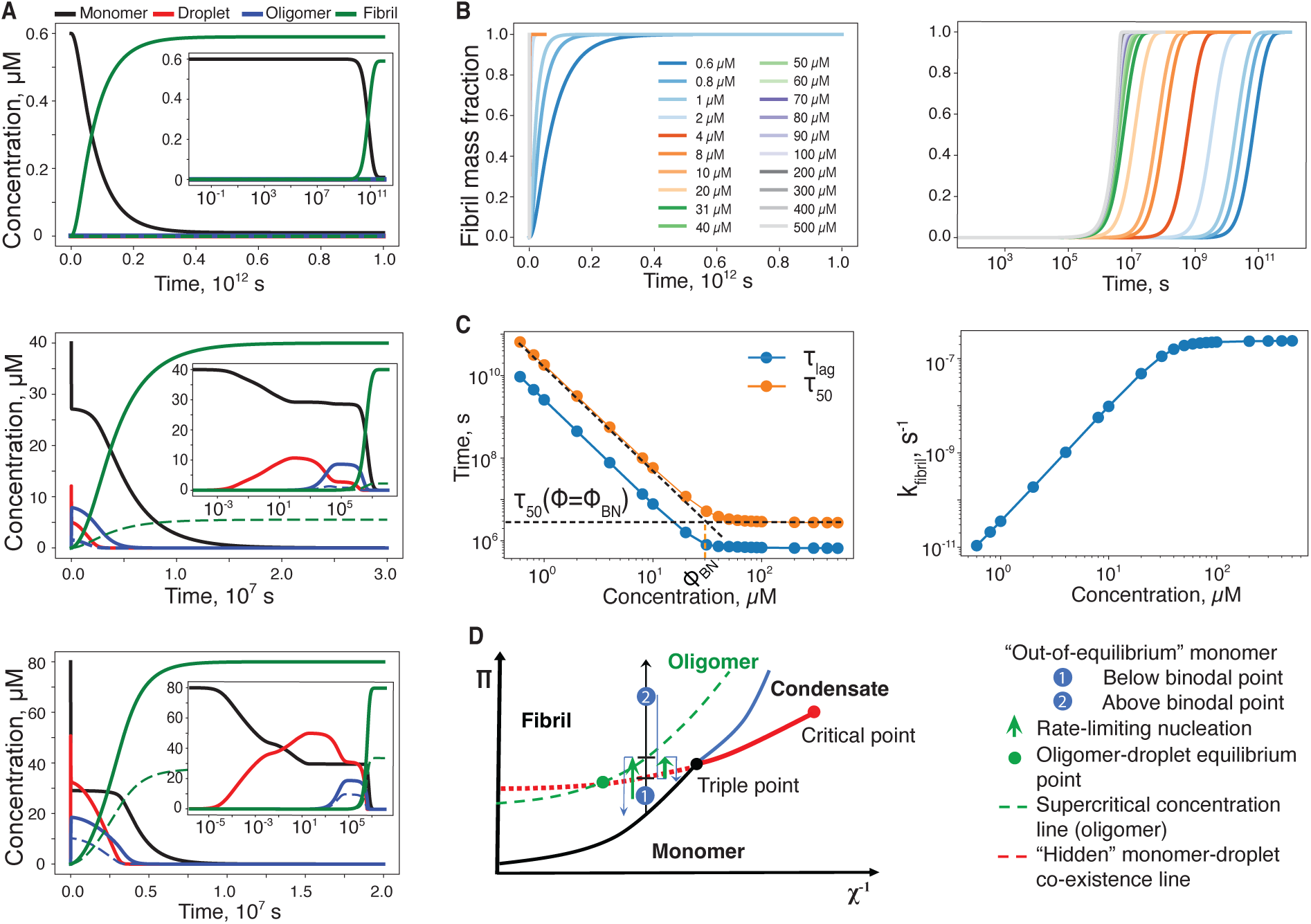
Aggregation kinetics of the global model. (**A**) Time evolution of total proteins in the form of monomers, droplets, oligomers, and fibrils at low (0.6 μM, top panel), intermediate (40 μM, middle panel) and relatively high (80 μM, bottom panel) concentrations with respect to the binodal concentration Φ_BN_ = 30 μM. The linear-log plot in the inset highlights the fast events, such as monomer-droplet equilibration and oligomerization. The dashed blue and green lines represent oligomers and fibrils nucleated out of the droplets. (**B**) Time evolution of the normalized fibril mass fraction of a wide range of concentrations shown in both linear (left) and linear-log (right) plots. (**C**) Fibrillization kinetic parameters including lag-time τ_lag_ and half-time τ_50_ (left) and fibril growth rate k_fibril_ (right) are shown as a function of total protein concentration. For the half-time plot (orange), the dashed back lines are fitting lines below and above Φ_BN_. (**D**) Phase diagram of amyloid protein states of monomers, droplets and fibrils similarly to the classical phase transitions between gas, liquid, solid states of matter. Here, the osmotic pressure π (proportion to the concentration of non-interacting monomers) and the inverse Flory-interaction parameter χ^−1^ are analagous to pressure and tempeature in the standard P-T phase diagram. The solid lines denote co-exisisting lines between different states, while the dashed lines denote transient co-existing lines between monomers and droplets and between monomers and oligomers within the fibril phase.

**Table 1.**
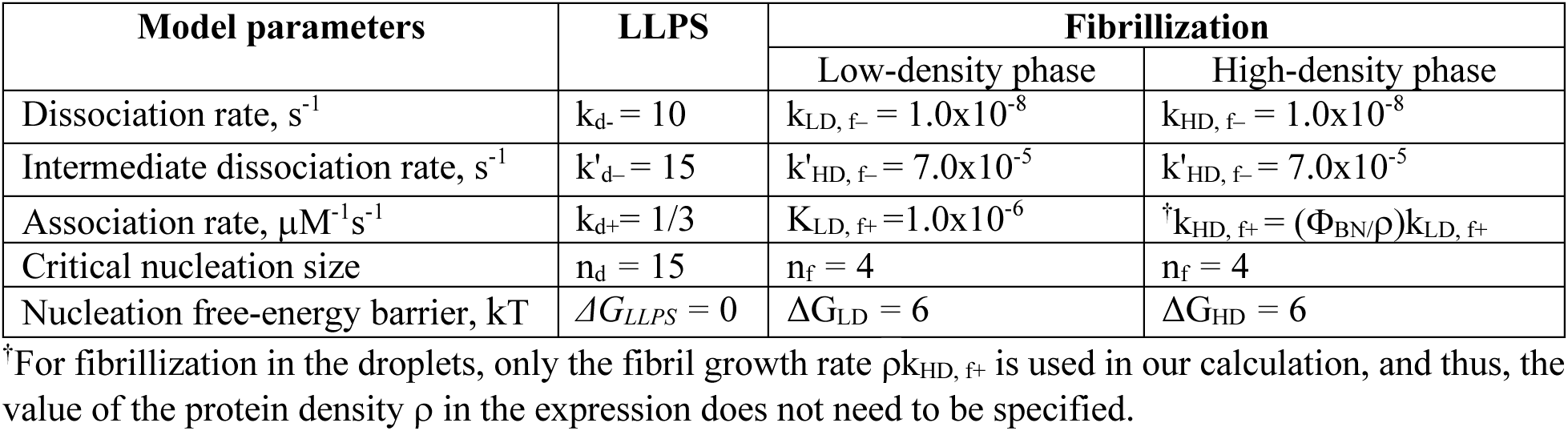
Model parameters for LLPS and for fibrillization in both high- and low-density liquid phases. The nucleation-dependent kinetics model (49) as shown in **Fig. 1C** is used. The threshold concentrations are Φ_CR_ = 10 nM, Φ_BN_ = 30 μM, and Φ_SU_ = 70 μM. We also set σ =1 and ΔΔG=0.

We first examined the aggregation kinetics for the total concentrations below, near and above Φ_BN_ (e.g., 0.6, 40, and 80 µM in **Fig. 2A**). At the low concentration regime below Φ_BN_, the droplet state was not stable and fibrillization occurred predominantly in the solution phase. Fibrils grew in the solution phase through monomer addition until the monomer-fibril equilibrium was reached. Throughout this aggregation process, both the droplets and oligomers were unstable with their populations being largely negligible. As expected, the droplet state became stable only near or above Φ_BN_ (e.g., the red lines in **Fig. 2A**). Droplets formed rapidly due to their fast association/dissociation rates (**Table 1**) and reached a transient monomer-droplet equilibrium before the formation of oligomers (blue lines) and fibrils (green lines). At the total protein concentration of 80 µM, there was an intermediary monomer-droplet plateau before the monomer concentration decreased to ~30 µM, corresponding to the formation of droplet intermediates that were included as part of the droplet phase. Oligomers formed out of both phases and underwent nucleated conformational conversion into fibril seeds, which then grew into fibrils (e.g., dashed lines corresponding to oligomers and fibrils originated from the droplets). With the fast monomer-droplet equilibrium, insofar as the droplet phase persisted, the monomer concentration in solution (i.e., the low-density liquid phase) remained approximately at the binodal concentration Φ_BN_, despite the constant net flux of proteins out of the solution phase to form oligomers and fibrils. Only where the droplets were fully depleted, the monomer concentration started decreasing until a monomer-fibril equilibrium was reached. As total protein concentration increased, more oligomers and fibrils were formed inside the droplets.

To better illustrate the concentration-dependence of fibrillization kinetics, we plotted the time evolution of normalized fibril mass fraction with different total protein concentrations in both linear and linear-log plots (**Fig. 2B**, shorter timescales for the linear plots shown in **Fig. S1**). At all concentrations, the fibrillization of the model system followed sigmoidal kinetics. At concentrations below Φ_BN_, the aggregation kinetics displayed a strong concentration-dependence showing steady and rapid decrease of aggregation times with increasing total protein concentrations. Above Φ_BN_, the aggregation kinetics saturated with increasing concentrations. The concentration-dependence of fibrillization kinetics can be further demonstrated by calculating the aggregation lag-time (τ_lag_), half-time (τ_50_) and fibrillization rate (k_fibril_) as a function of total protein concentration (**Fig. 2C**). Here, we followed the standard approaches (62, 63) in the literature to compute these kinetics parameters. Specifically, the half-time was defined as the time required to reach 50% of the maximum fibril mass. The lag-time and growth rates were estimated by drawing a tangent line of the fibrillization curve at the half-time, where the intersection with the time-axis marked τ_lag_ and the slope corresponding to k_fibril_. Indeed, below Φ_BN_, all three kinetics parameters followed a power-law scaling with the total protein concentrations (e.g., dashed linear fitting line in the log-log plots in **Fig. 2C**), where τ_lag_ and τ_50_ decreased and k_fibril_ increased with the increasing protein concentration. Above Φ_BN_, each of the three kinetics parameters approached a constant with increasing concentrations.

The power-law scaling at low concentrations is consistent with the classical nucleation theory. Using the same approximation of rapid pre-equilibrium of monomers with weakly populated oligomers as in Powers and Powers (48), an analytic expression of the half-time can be obtained (detailed derivation in the **SI**):

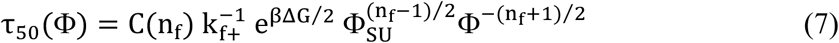

where C(n) = sech^−1^[2^−(n+1)/2^] is a constant and Φ the total protein concentration. Compared to Powers and Powers (48), the extra term of e^βΔG/2^ resulted from the conformational conversion barrier Δ*G* for fibrillization as introduced by Serrano et al.(49) (e.g., **Fig. 1C**). At high concentrations, saturation occured because the increase of protein concentrations above Φ_BN_ only changed the partition of proteins between the co-existing phases with fixed protein concentrations (i.e., Φ_BN_ and ρ, the upper and lower binodal points along the tie-line in **Fig. 1B**). As discussed above, aggregation inside the droplets was thermodynamically equivalent to that in the co-existing monomer solution. In this model system (**Table 1**), we also assumed that the aggregation kinetics were the same also with σ =1 and ΔΔG=0. As a result, the aggregation kinetics became saturated with increasing concentrations above Φ_BN_, and the half-time τ_50_ approached a constant of approximately τ(Φ = Φ_BN_) in Eq. 7 (the horizontal fitting line in **Fig. 2C**).

The above results may also be understood using the proposed phase diagram of amyloid protein monomer, droplet, and fibril states that has been proposed in analogous to the phase diagram of gas, liquid and solid states of matter (54). Here, we introduced a supercritical concentration line Φ_SU_(χ) – i.e., the transient monomer-oligomer co-existing line – within the fibril phase in an updated phase diagram (the green line in **Fig. 2D**). Both droplets and oligomers were intermediate states during fibrillization: their chemical potentials depended on the solvent condition and the protein solute (**Fig. 1E**), and their relative stabilities might swap at high enough χ-values (e.g., the line intersect highlighted as the green circle in **Fig. 2D**). Our parameter selection (**Table 1**) corresponded to the protein-solvent condition between the oligomer-droplet equilibrium point and the triple point. Amyloid aggregation can be viewed as an equilibration process from an initially homogeneous, out-of-equilibrium monomer solution toward thermodynamically stable fibrils via oligomerization, nucleated conformational conversion and growth – e.g., the vertical relaxation path from a point within the fibril phase back to the fibril phase boundary via the obligatory formation of oligomers (**Fig. 2D**). For a starting point below the monomer-droplet co-existing line (i.e., concentrations below Φ_BN_), the rate-limiting step of fibrillization is the transition towards oligomers, where the energetic separation deceases with increasing total protein concentrations. At initial concentrations above the monomer-droplet co-existence line, the system rapidly relaxes back to this line, after which the rate-limiting step for oligomer formation becomes constant, leading to saturation of the aggregation kinetics.

### Dependence of fibrillization on different model parameters

We first examined the impact of LLPS on fibrillization by comparing the aggregation kinetics with and without LLPS. For the cases without LLPS, we simply turned off the formation of droplets in solving Eqs. 1–6. As expected, the fibrillization kinetics was influenced by LLPS only above the binodal point, displaying large deviations for two of the cases (e.g., the solid and dashed lines in yellow at 80 μM and in black at 400 μM in **Fig. 3A**). Compared to the saturated aggregation kinetics with droplet formation above Φ_BN_, without LLPS the aggregation lag-time kept decreasing at a slower rate with increasing total protein concentration above Φ_SU_. Such differences can be better illustrated in the log-log plot of τ_50_ as a function of total protein concentration (blue line in **Fig. 3B**). Without LLPS, the decrease of τ_50_ with increasing protein concentration slowed near and above the supercritical concentration, Φ_BN_, as reported by previous models (48, 49). Above the supercritical concentration, the oligomers were more stable than monomers and the fibrillization rate was dictated by the concentration of oligomers, resulting in a power-law scaling of 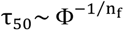 (the black dashed fitting line in **Fig. 3B**, details of the derivation in the **SI**).

**Figure 3.**
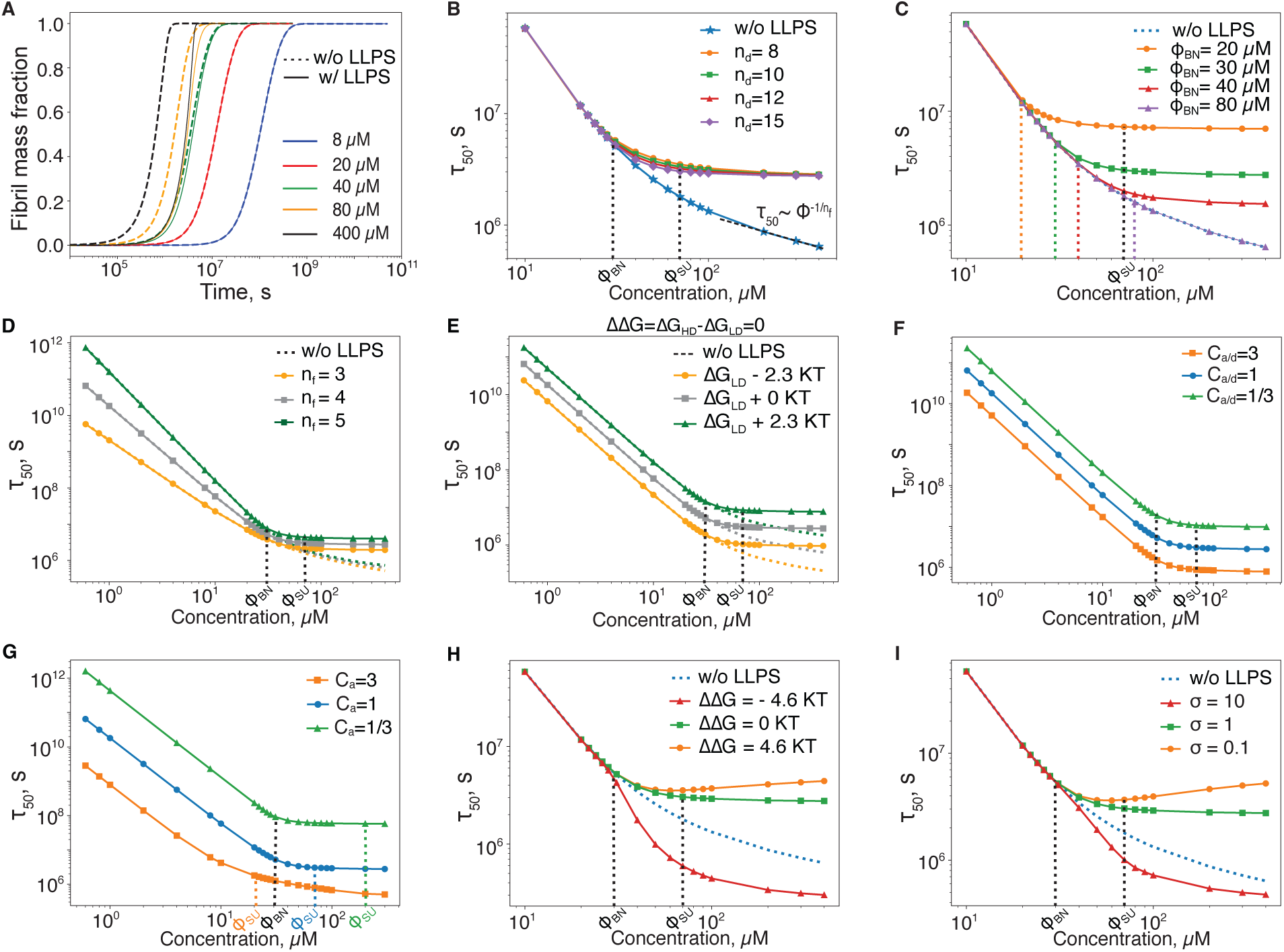
Dependence of fibrillization on model parameters. (**A**) Time evolution of the normalized fibril mass fraction for different monomer concentrations with (solid lines) and without (dashed lines) LLPS. Half-time τ_50_ versus concentration curve for varying (**B**) Critical size of the droplet n_d_, (**C**) Binodal point Φ_BN_, (**D**) Critical nucleus size n_f_ of fibrillization, (**E**) The oligomer conformational conversion barrier ΔG in both protein phases by keeping ΔΔG=0, (**F**) All fibrillization association/dissociation rates in both phases by the rescaling coefficient C_a/d_ such that supercritical and critical concentrations (Φ_SU_, Φ_CR_) are maintained, (**G**) Only the fibril association rate in both phases by the rescaling coefficient C_a_ such that both supercritical and critical concentrations are changed accordingly, (**H**) ΔΔG that governs the difference of oligomer conformational conversion barriers in two phases, and (**I**) Coefficient σ that determines the difference of fibrillization rates in the two phases.

The different asymptotic behaviors for droplet-induced saturation and oligomer-induced slowing down of fibrillization at high protein concentrations can be understood by their different size limits – droplet sizes range from n_d+1_ to infinity, but oligomer sizes vary from 2 to n_f_. For LLPS, increasing total protein concentration above the binodal concentration Φ_BN_ only led to more proteins partitioned to the droplet phase with the co-existing monomer concentration staying constant. During oligomerization, increasing total protein concentration above the supercritical concentration Φ_SU,_ on the other hand, led to continuous concentration increases of both oligomers and monomers, only at a slower pace.

We next assessed different model parameters on fibrillization kinetics over a wide range of protein concentrations. We first focused on parameters associated with LLPS. The droplet nucleation size, n_d_, determines the cooperativity of droplet formation. By adjusting n_d_ while keeping other parameters the same as in **Table 1**, our results showed that higher n_d_ led to faster aggregation kinetics with shorter τ_50,_ although the difference appears mainly around Φ_BN_ (**Fig. 3B**). Systems with a large n_d_ required the cooperative assembly of more monomers, resulting in fewer total droplets and a higher concentration of co-existing monomers (**Fig. S2**). Hence, our result suggested that the fibrillization kinetics of the monomer-droplet mixture was mainly determined by the concentration of the co-existing monomer solution. Because droplet formation is typically orders-of-magnitude faster than fibrillization, we found that changes in the LLPS rates without changing Φ_BN_ did not affect amyloid aggregation at different concentrations. Varying the binodal concentration Φ_BN_, on the other hand, significantly impacted amyloid aggregation by changing the monomer-droplet equilibrium, thus changing the droplet-induced saturation of amyloid aggregation (**Fig. 3C**). The lower the Φ_BN,_ the slower the saturated fibrillization kinetics. Interestingly, when Φ_BN_ was above the supersaturation concentration Φ_SU,_ the fibrillization kinetics overlapped with those without LLPS at different concentrations (e.g., the purple curve in **Fig. 3C**). When Φ_BN_ > Φ_SU_, oligomers were thermodynamically more stable than droplets, and thus the initially formed droplets above Φ_BN_ were rapidly converted into oligomers, which in turn determined the fibrillization kinetics (**Fig. S3**).

In terms of fibrillization-related parameters, we first kept σ=1 and ΔΔG=0 (i.e., fibrillization in the two phases were both thermodynamically and kinetically equivalent to each other) while adjusting other parameters applied to fibrillization both in the solution and droplets (**Fig. 3D-G**). By varying fibril nuclei size n_f_ alone, our results showed that systems with larger n_f_ values featured a sharper decrease in τ_50_ with increasing protein concentration and had slower aggregation kinetics both below and above Φ_BN_ (**Fig. 3D**). As shown in **Fig. S4A**, oligomers were less populated for larger n_f_. The results on varied n_f_ were fully consistent with Eq. 7. As expected from the same equation, increasing nucleated conformational conversion barrier ΔG in both protein phases led to the slowing of aggregation kinetics with increased τ_50_ (**Fig. 3E**). With a greater ΔG, oligomers took a longer time to form fibril seeds (**Fig. S4B)**. By uniformly adjusting all fibrillization rates with a rescaling coefficient C_a/d_ that had Φ_SU_ unmodified, higher rates with larger coefficients gave rise to faster kinetics and smaller τ_50_ (**Fig. 3F**), consistent with the prediction from Eq. 7 with τ_50_~C_a/d_^−1^. Adjusting either association or dissociation rates of fibrillization could modify Φ_SU_. In this case, we simply scaled the fibril association rate alone with a coefficient C_a_. The increase of fibril association rate would result in a decrease of Φ_SU_, and τ_50_ became proportional to C_a_^−(n+1)/2^, as predicted by Eq. 7. Indeed, large C_a_ led to faster aggregation kinetics with shorter τ_50_ (**Fig. 3G**). When Φ_SU_ > Φ_BN_, the droplet-induced saturation of aggregation kinetics was observed at concentrations above Φ_BN_. As Φ_SU_ decreased below Φ_BN_, the supersaturation-induced slowing of fibrillization kinetics was observed around and above Φ_SU_.

With ΔΔG ≠ 0 or σ ≠ 1, fibrillization in droplets was kinetically different from that in the solution. Varying ΔΔG (**Fig. 3H**) and σ (**Fig. 3I**) mostly affected the fibrillization above Φ_BN_ where droplets were stable. Above Φ_BN_, increasing total protein concentration led to more proteins partitioned into the droplet phase, and the overall fibrillization kinetics of the droplet-monomer mixture subsequently became more dominated by that in the droplets. Under conditions σ < 1 and/or ΔΔG > 0, fibrillization was slower in the droplets than in the solution, and thus, the overall fibrillization kinetics slowed (e.g., τ_50_ increased) with increasing total protein concentration (**Fig. 3H,I** and **Fig. S4C,D**). With σ > 1 and/or ΔΔG < 0, fibrillization in the droplets was faster compared to that in the solution, and the overall fibrillization kinetics accelerated (e.g., τ_50_ decreased) with increasing total protein concentration.

### Delineating the effects of droplet formation on amyloid aggregation

To study the effects of LLPS on fibrillization, typical experimental approaches to facilitate droplet formation include the addition of multivalent scaffolding agents (32, 41), crowders (32, 64), or adjusting solution conditions such as pH (32, 65) and salt concentration (41, 45, 65). These interventions are intended to modulate the phase boundary of the condensate in the phase diagram (**Fig. 2D**), subsequently affecting fibrillization kinetics. Interestingly, seemingly contradicting phenomena including fibrillization-promotion (66–68), inhibition (42, 44, 69) and even biphasic effects (45, 65) have been documented in the literature. Here, we categorized different LLPS-modulating approaches into those that mainly impact either the monomer solution or condensate phases, and those that affect both phases, and applied our global model to delineate their corresponding effects on fibrillization.

#### Impacting the low-density liquid phase

Many experimental studies have shown that crowding agents such as polyethylene glycol, dextran, and Ficoll can promote LLPS and fibrillization (32, 66, 67). These crowding agents increase the effective protein concentration by exerting steric hinderance to the monomers in the solution. Upon LLPS, crowders stay in the protein monomer solution instead of the droplets. Assuming no binding interactions with proteins and their assemblies (i.e., no impact on the dissociation rates), the crowder effect was modeled by modulating the association rates for droplet formation k_d+_ and fibrillization in the solution k_LD, f+_ with a rescaling coefficient C_sol_ > 1, while keeping other parameters unchanged (see **Table 1**). Here, the fibril association rate k_HD, f+_ was not rescaled because crowders were not partitioned in the droplets during LLPS, and thus, effectively σ = C_sol_^−1^ after rescaling. Solving Eqs. 1–6 with C_sol_ = 3 for different total protein concentrations, our results showed that crowders indeed accelerated fibrillization with reduced τ_50_ compared to the unmodulated system, at low concentrations and even after saturation of fibrillization kinetics due to the lowered binodal concentration Φ_BN_ until saturation in the reference system was also reached (**Fig. 4A**). Interestingly, rescaling with a coefficient C_sol_ = 1/3 can be used to model the protein-sequester effect (69–71) of certain biomolecules that bind amyloid proteins in the solution but not in the droplets, which can inhibit or delay fibrillization with an opposite effect to crowders (**Fig. 4A**).

**Figure 4.**
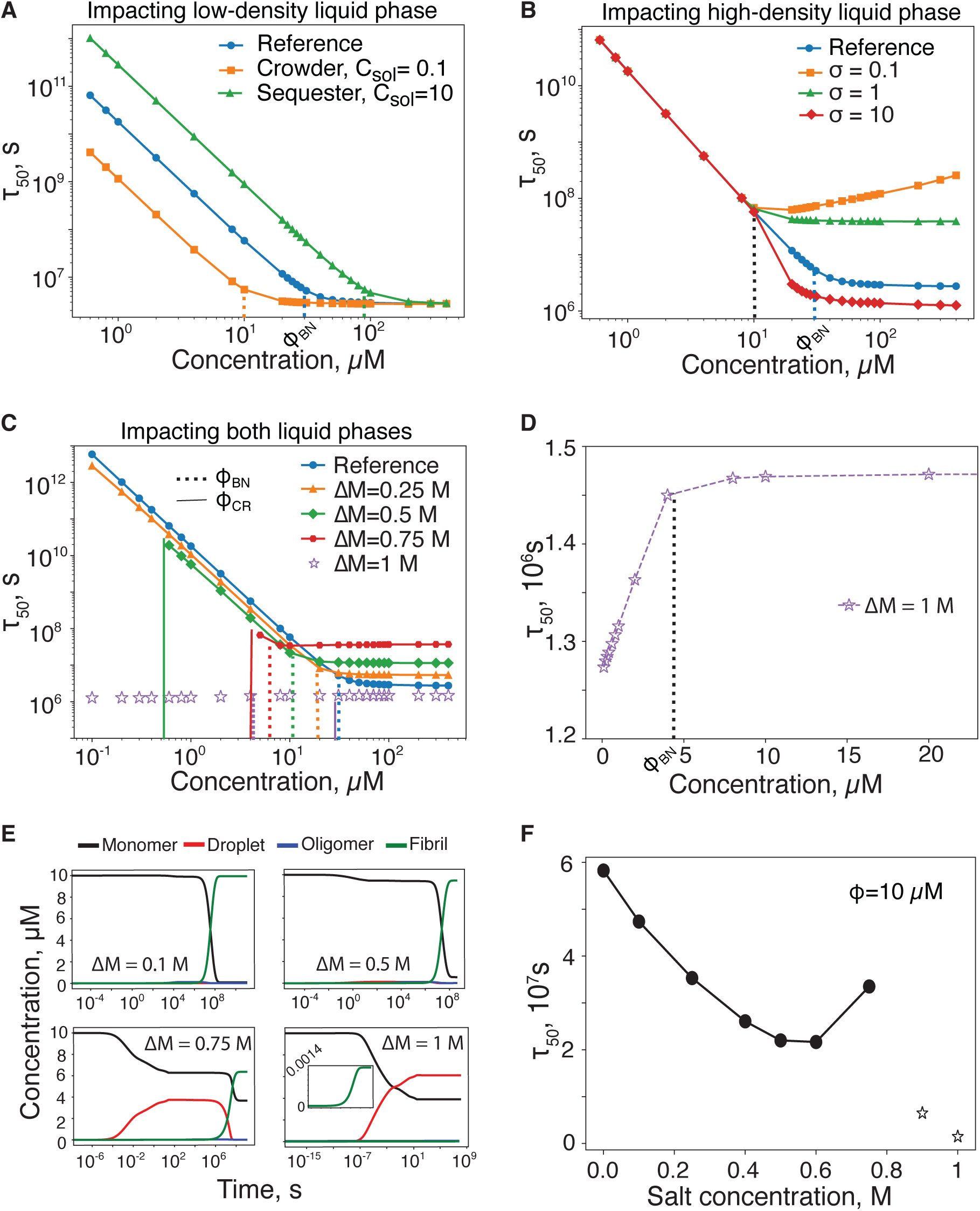
Modulation of fibrillization kinetics by different droplet-promoting approaches. (**A**) Impacting proteins in the solution phase by either molecular crowding or monomer-sequestering agents. The effects were modeled by scaling the droplet association rate and the fibril association rate in the solution, C_sol_. Crowders (C_sol_ = 1/3) promote fibril formation by enhancing effective monomer concentration while monomer-sequestering molecules (C_sol_ = 3) delay fibrillization by lowering effective monomer concentration. (**B**) Impacting proteins in the droplets by multivalent scaffolding proteins, which lower the co-existing monomer concentrations but may promote (σ > 1) or delay (σ < 1) the fibrillization above the binodal point depending on their interactions with monomers in the droplet. (**C**) Impacting proteins in both phases, salt displays intricate effects on fibrillization. The salt-induced screening effect on electrostatic interactions was modeled by modulating both LLPS and fibrillization rates with the approximation and parameters listed in **Table 2**. (**D**) Half-time τ_50_ vs protein concentration at a high salt concentration of 1 M, where fibrils are less stable than droplets, Φ_BN_ < Φ_CR_. (**E**) Time evolution of monomer, droplet, oligomer and fibril mass concentrations for 10 µM total proteins at different salt concentrations. (**F**) Non-monotonic or biphasic concentration-dependent effect of droplet-promoting salt on the fibrillization of 10 µM total proteins. Open stars represent cases where fibrils are not stable compared to droplets.

**Table 2.** List of parameters used to model salt-induced rate changes. Under changes to salt concentration ΔM with respect to the reference model, each rate was modulated according to the linear energy approximation k = k_ref_ exp(C_Salt_ΔM) due to the screening effect of salts on electrostatic interactions. Here, k_ref_ is the reference rate in Table 1, and C_Salt_ is the salt-destabilizing coefficient listed below in the unit of M^−1^. Since salt affects protein interactions equally in both the solution and droplets, we kept σ =1 and ΔΔG=0 as in Table 1.

|  | LLPS | Fibrillization |  |
| --- | --- | --- | --- |
|  |  | Low-density phase | High-density phase |
| Dissociation rate | 0.0 | 10.0 | 10.0 |
| Intermediate dissociation rate | 0.0 | 2.0 | 2.0 |
| Associate rate | 2.0 | 2.0 | 2.0 |

#### Impacting the high-density liquid phase

Some recent studies have shown that promoting droplet formation of amyloid proteins by multivalent binding molecules such as nucleotides, chaperones, or scaffold proteins could prevent or delay fibril formation (42, 45, 72–75). These molecules are mostly partitioned in the phase-separated droplets together with amyloid proteins. Hence, for these droplet-promoting co-condensates, the effect can be modeled by rescaling the droplet dissociation rate k_d-_ with a coefficient C_drop_ < 1, leading to a lower Φ_BN_. Due to heterotypical interactions between amyloid protein and the co-solutes, fibrillization in the droplets can be kinetically distinct from those in solution. For the monomer-droplet mixture with a lower co-existing monomer concentration, the overall fibrillization kinetics was slower with a longer half-time τ_50_ compared to the homotopical system where σ ≤ 1 (e.g., σ = 1, 0.1 **Fig. 4B**), consistent with prior experimental reports (42, 72, 74). However, under special cases where the binding of amyloid proteins with co-condensates promoted fibrillization and leads to faster fibrillization in the droplets than in the solution with σ > 1, overall fibrillization could also be accelerated compared to the reference system (e.g., σ = 10 in **Fig. 4B**). For instance, the reported acceleration of IAPP by a scaffold star-polymer with rod-like arms to template fibril formation can be explained by the latter case (76).

#### Impacting both low and high-density liquid phases

Multiple studies have reported that increasing salt concentrations can promote both LLPS and fibrillization of different amyloid proteins (68, 77, 78). By studying the salt effects over a wide concentration range on both LLPS and fibrillization of α-synuclein, Sternke-Hoffmann et al. (45) recently showed that increasing salt concentration promoted LLPS monotonously, but displayed a biphasic effect on fibrillization where the initial fibrillization-promotion effect at low to moderate salt concentrations was reversed above a certain threshold. At very high salt concentrations, they observed a remarkable dissolution of preformed fibrils into droplets after prolonged incubation. It is well known that salt promotes droplet formation by reducing electrostatic interactions via screening. Upon LLPS, salt ions are present in both phases and exert a screening effect equally on proteins partitioned into the two phases.

α-synuclein protein possess a highly negative net charge of −12e at physiological pH, and thus, the presence of salt ions reduces electrostatic repulsions upon the self-association of α-synuclein in both LLPS and fibrillization. Salt can also weaken salt-bridges and hydrogen bonds that stabilize the aggregated forms of droplets, oligomers and fibrils, thereby increasing their corresponding dissociation rates. Assuming a linear concentration dependence of salt-induced changes on interaction potentials, salt-induced increase of different rates can be approximated by rescaling with exp(C_Salt_ΔM), where ΔM denotes the change of salt concentration with respect to the reference system and salt-screening coefficient C_Salt_ is proportional to the number of relevant electrostatic interactions (all C_Salt_ values listed in **Table 2**). Since the association rates for both LLPS and fibrillization are related to monomer binding, we assumed the same salt-destabilizing coefficient for k_d+_, k_LD,f+_ and k_HD,f+_. Among different aggregates, the droplets entail the weakest intermolecular interactions (**Fig. 1D,E**), and we assigned C_Salt_ = 0 to the droplet dissociation rate so that the binodal concentration Φ_BN_, defined as k_d–_/k_d+_, decreased with increasing salt concentration (i.e., droplets became more stable). Fibrils are stabilized by extensive intermolecular hydrogen bonds while oligomers possess less intermolecular interactions with transient hydrogen bonds. Hence, we assigned a higher C_Salt_ value for fibril-dissociation rates, k_LD,d–_ & k_HD,d–_, than the oligomer-dissociation rates, k’_LD,d–_ & k’_HD,d–_. For simplicity, we chose the same C_Salt_ value for oligomer-dissociation rates as for the fibril association rates, such that with increasing salt concentrations the supercritical point Φ_SU_ remained same but the critical concentration Φ_CR_ increased (i.e., fibrils became less stable). Hence, salt increases led to a shift of boundaries or co-existing lines in the phase diagram (**Fig. S5)**, subsequently modulating fibrillization kinetics.

Using the salt-dependent model, we computed fibrillization half-time over a wide range of protein concentrations for different salt concentrations (**Fig. 4C,D**). With increasing salt concentrations, increased fibril association rate led to *accelerated fibrillization* kinetics with reduced τ_50_ at low concentrations prior to droplet-induced saturation. The decreased Φ_BN_ with increasing salt concentrations (highlighted as dashed vertical lines in **Fig. 4C**), on the other hand, led to *delayed fibrillization* kinetics with increased τ_50_ in the saturated stage at high protein concentrations above the corresponding binodal point. These two competing effects led to the experimentally reported salt-induced biphasic effects on α-synuclein fibrillization by Sternke-Hoffmann et. al.(45) – e.g., the fibrillization half-time at a fixed protein concentration Φ = 10 μM with increasing salt concentration as in our model (**Fig. 4E,F**).

Additionally, increasing salt concentration also led to destabilization of fibrils with increasing critical concentration Φ_CR_ (highlighted as solid vertical lines in **Fig. 4C**), below which fibrils were unstable. Especially when Φ_CR_ > Φ_BN_ (e.g., at 1M salt, **Fig. 4C,D**), the fibrils were unstable even with the total protein concentration above the critical concentration because of the readily formed droplet-monomer mixture where the effective monomer concentration stayed at Φ_BN_. At concentrations below Φ_CR_, the amounts of formed fibrils and oligomers were very little with the monomer concentration Φ_m_ nearly unperturbed, and there, the kinetics followed a simple exponential growth, ~ 1 - exp[-k_f+_t(Φ_m_ - Φ_CR_)] (e.g., as illustrated by the fibrillization time-evolution of 10 μM total proteins at 1M salt in **Fig. 4E**). With little conversion of monomers into oligomers and fibrils, monomer concentration Φ_m_ effectively equaled to the total concentration Φ, and thus, the half-time depended on Φ according to, τ_50_ ~1/(Φ-Φ_CR_), below Φ_BN_, but saturated above Φ_BN_ (**Fig. 4D**). This result explained the reported disaggregation of α-synuclein fibrils into droplets at very high salt concentrations (45).

Taken together, our analyses suggest that while various droplet-promoting approaches all stabilize the droplet state and lower the monomer-droplet co-existing line in the phase diagram, they can also differentially modulate the kinetic and thermodynamic properties of other states – monomers, oligomers, and fibrils – as illustrated in the SI for crowders (**Fig. S6**), co-condensates (**Fig. S7**), and salts (**Fig. S5**). These modulations, in turn, can lead to different outcomes to the fibrillization kinetics and thermodynamics in the low protein concentration regime below the binodal point and in the high protein concentration regime above the binodal point. Our global thermodynamic-kinetic model thus resolves the diverse and confounding effects of LLPS on protein fibrillization reported in the literature.

### The peculiar cases of slowed fibrillization with increasing protein concentration

As illustrated by numerous prior experimental studies (48, 49) and predicted by the classical nucleation theories (48, 49) and their variations (51), increasing protein concentrations accelerate fibrillization, though the kinetics can slow down at high concentrations. However, it has been reported that human calcitonin (hCT) exhibited a non-canonical delaying of aggregation kinetics (i.e., increase of half-times) (55) with increasing protein concentrations. Interestingly, our model captured this peculiar fibrillization behavior at concentrations above the binodal point with ΔΔG > 0 (**Fig. 3H**) or σ < 1 (**Fig. 3I**) – i.e., fibrillization in the droplets has either a higher nucleated conformational conversion barrier or a slower association rate. To further demonstrate this non-canonical behavior, we illustrated the normalized fibril growth above the binodal points with σ = 0.1 (**Fig. 5A**) and ΔΔG = 4.6 KT at different protein concentrations (**Fig. 5B**). Hence, our model offered a plausible mechanism for understanding the experimentally-observed slowing of hCT fibrillization with increasing protein concentrations (55).

**Figure 5.**
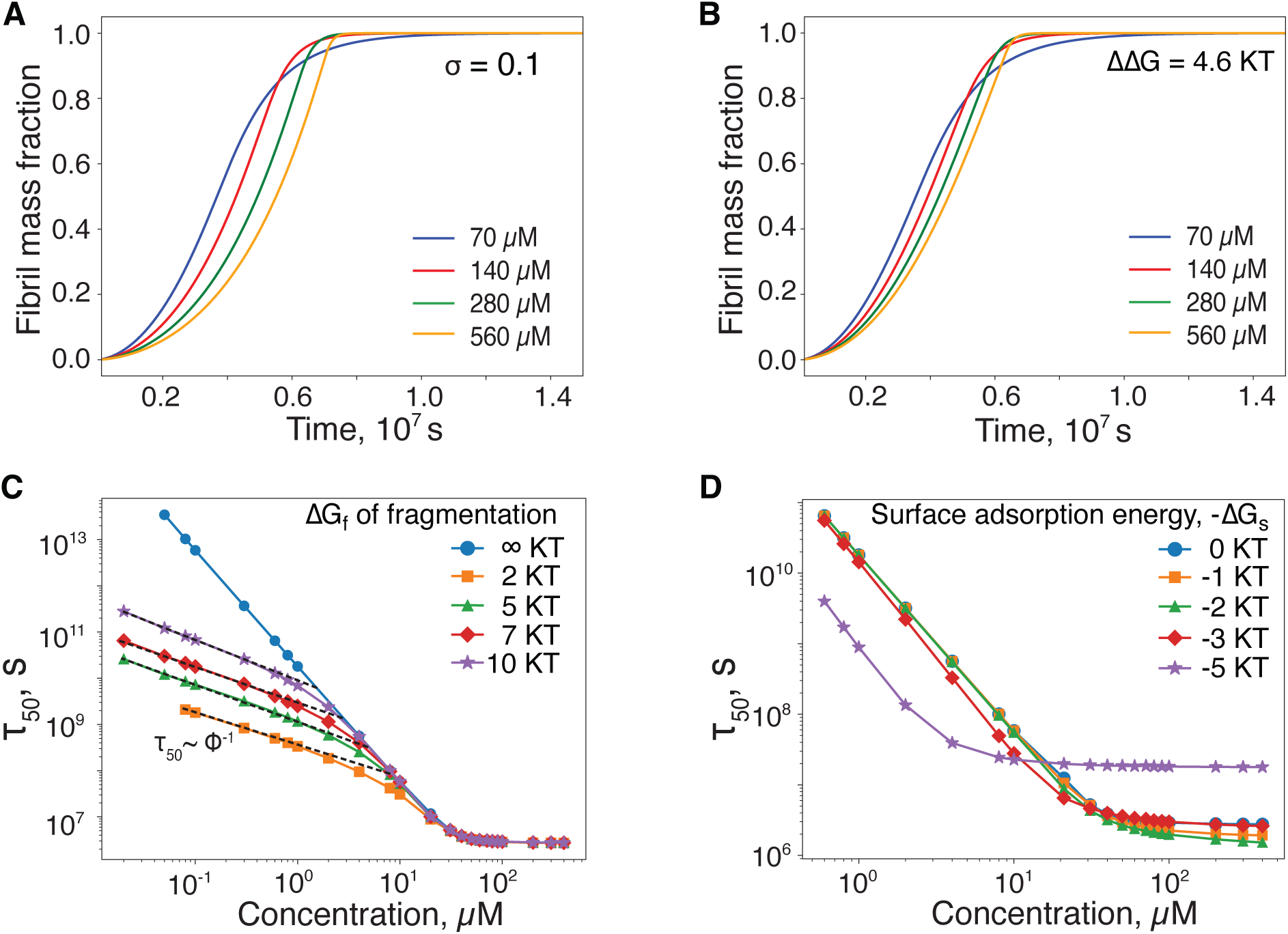
Aggregation kinetics of peptides under special cases. Under the conditions of (**A**) the coefficient σ = 0.1 and (**B**) ΔΔG = 2 ln(0.1) KT = 4.6 KT, the fibrillization kinetics slowed with increasing protein concentrations. (**C**) Effect of fragmentation on fibril formation kinetics with fibrillization half-time τ_50_ shown as a function of total protein concentration for different fragmentation rates. ΔG_f_ determines how much fibril fragmentation rate is slower than the fibril dissociation rate. ΔG_f_ = ∞ KT corresponds to the reference system without fragmentation. (**D**) Effect of secondary nucleation on fibril formation kinetics with τ_50_ shown as a function of total protein concentration for different extent of protein enrichment on the surface of pre-formed fibrils, quantified by ΔG_s_ – the protein-binding energy on the fibril surface. ΔG_s_ *=* 0 KT denotes the reference system without secondary nucleation.

### Effects of secondary nucleation and fragmentation in the present model

Fibril-dependent fragmentation and secondary nucleation have been shown to play important roles in amyloid aggregation (15, 49, 51, 79, 80). Here, we also examined the effects of these two processes on amyloid aggregation within the context of our new model. Amyloid fragmentation involves breaking up of fibrils into small fragments, which in turn increases the active sites for fibril growth via monomer addition at the fibrillar tips. Compared to monomeric fibril dissociation at fibril ends where terminal monomers are only partially ordered, fragmentation occurring within the fibril core where the structure is more ordered is energetically unfavorable, and thus, less frequent. To account for this difference, we introduced an energy term, ΔG_f_ of fragmentation, which quantifies how much lower the fragmentation rate k_fragment_ is compared to the monomeric fibril dissociation rate, k_fragment_ = k_f-_ exp(-βΔG_f_). Fragmentation elicited a strong effect at low protein concentrations, significantly reducing half-time τ_50_ (**Fig. 5C**) and lag-time τ_lag_ (**Fig. S8A**). Under this low concentration condition regime where the fragmentation effect dominated, the half-time τ_50_ followed a different power-law scaling with concentration, 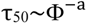 (with a ≈ 1, fitted by black dashed lines, **Fig. 5C**). The larger ΔG_f_, the lower threshold concentration under which the power-law scaling deviated from the reference case without fragment (i.e., ΔG_f_ = ∞).

Secondary nucleation involves the formation of new fibril seeds through the adsorption of monomers onto the surfaces of preformed fibrils. We introduced a surface adsorption energy, −ΔG_s_, between amyloid proteins and fibril surfaces, which resulted in the enrichment of local protein concentration on the fibril surface by the factor of exp(β ΔG_s_), thereby facilitating secondary nucleation both in the solution and droplets (details in the **SI**). For simplicity, we assumed other fibrillization parameters were unchanged on the fibril surface. As the nucleation rate of fibril seeds was proportional to the n_f_’th-power of the protein concentration, the seeding rate in secondary nucleation was greatly amplified by a factor of exp(n_f_ βΔG_s_) due to local concentration increases. With a relatively weak ΔG_s_ (e.g., 1 & 2 KT in **Fig. 5D**), secondary nucleation did not significantly affect half-time τ_50_ (**Fig. 5D**) and lag-time τ_lag_ (**Fig. S8B**) at low concentrations compared to the reference system without secondary nucleation. Only at higher concentrations, secondary nucleation showed a significant reduction in τ _50_, suggesting its increased relevance as the total protein concentration increased. As the surface adsorption energy increased further (e.g., ΔG_s_ = 3KT and 5KT in **Fig. 5D**), secondary nucleation started dominating over primary nucleation, leading to self-amplifying at low concentrations below the binodal point. On the other hand, preformed fibrils in the droplets effectively served as a co-condensate to enhance droplet formation but with delayed fibrillization in the monomer-droplet mixtures (e.g., **Fig. 4B**).

## Conclusion

In conclusion, we have developed a global thermodynamic-kinetic model bearing the hallmarks of both LLPS and amyloid fibrillization. Specifically, we regarded protein condensates as a new state of amyloid protein solution in addition to the non-interacting monomer solution and fibrils, and oligomers as fibrillization intermediates. Different from traditional fibrillization models that treat the protein solution as homogenous, we considered fibril nucleation and growth in a heterogeneous monomer-droplet coexisting mixture. Based on the thermodynamics laws, fibrillization in the condensates is thermodynamically but not kinetically equivalent to that in the co-existing protein-poor solution. We adopted the classical nucleation theory to model the phase transitions of both LLPS and fibrillization and introduced parameters like σ and ΔΔG to describe the potential kinetic differences of fibrillization in solution from droplets. Together, our model recapitulated the power-law scaling of fibrillization kinetics at low protein concentrations as well as saturation of fibrillization kinetics at high concentrations due to the rapid formation of the monomer-droplet co-existing mixture of LLPS.

Critically, this global model allowed us to rationalize confounding reports about the effects of droplet formation on fibrillization, including fibrillization promotion, inhibition and even biphasic effects (42–45). By categorizing common droplet-promoting approaches used in experiments into those that impact either the solution monomers (e.g., crowders), droplets (e.g., co-condensates), or both (e.g., pH and salts), we showed that these approaches also affected other states while stabilizing the droplets (**Fig. S5-7**). By considering all causative factors in the model parameters, we reproduced all experimentally observed effects of droplets formation on LLPS, demonstrating a remarkable predictive power of the global model in encompassing the biophysical-biochemical processes of both LLPS and fibrillization. These results are physiologically relevant because cellular environments are highly complex with high density distributions of proteins and nucleotides that can serve as crowders, monomer-sequesters, or multivalent co-condensates, and also possess varying salt concentrations and pH values across cellular compartments or during various biological processes, such as the release of IAPP from the inter- to intracellular space of pancreatic beta cells or the splitting of one mother cell into two daughter cells in mitosis aided by the polymerization of microtubules and their associated protein tau. In addition, our results point to general mitigating amyloidosis strategies of delaying amyloid protein aggregation by monomeric-sequestering in solution (**Fig. 4A**), LLPS-promoting with co-condensates that do not reduce the free-energy barrier of nucleation and fibril growth (**Fig. 4B**), or effectively increasing the free-energy barriers associated with either nucleated conformation conversion or fibril growth (**Fig. 3E-G**).

In addition to the above-described applications, we have demonstrated the applicability of the model to the peculiar case of hCT amyloid aggregation, where retarded fibrillization was rendered with increasing protein concentration. Moreover, we have examined the effects of secondary nucleation and fragmentation in the context of our model. Together, this global thermodynamic-kinetic model can closely represent the behaviors of various protein-solvent systems and is applicable to diverse fibrillization pathways. The tunable parameters can also be used to validate approaches modulating amyloid aggregation. Therefore, this unified model of LLPS and amyloid aggregation represents a major advancement in the field of amyloidosis and is expected to facilitate our understanding and mitigation of the complex aggregation processes underscoring the pathophysiology of a range of neurodegenerative, systemic and metabolic disorders.

## Acknowledgements

This work was supported by NIH R35GM145409 and the Research Program of the South Carolina Alzheimer’s Disease Research Center. Computer simulations were run on the Palmetto high performance cluster at Clemson University and were supported by the multiscale computational modeling core of NIH P20GM121342. The content is solely the responsibility of the authors and does not necessarily represent the official views of the NIH.

## Data, Materials, and Software Availability

All data with corresponding python notebooks are available on Zenodo (https://doi.org/10.5281/zenodo.15499212) and the standalone Python scripts is available on GitHub (https://github.com/kamalbhandari/AMYLOID_LLPS).

## Author Contributions

F.D. conceived the project; K.B. and F.D. developed the models, performed calculations and data analysis; K.B., Y.S., H.T., P.C.K. and F.D. wrote the manuscript. All authors provided inputs and agreed on the presentation of the manuscript.

