## Supplementary Information for "A Global Thermodynamic-Kinetic Model Capturing the Hallmarks of Liquid-Liquid Phase Separation and Amyloid Aggregation"

### Supplementary Methods and Discussions

#### A. Derivation of the rate-mass equations

We adopted an approach similar to that used by Serrano et al.(1) to model the phase transitions of both LLPS and fibrillization, where a pre-equilibrium approximation was applied to smaller intermediates and multiple monomers were assumed to assemble simultaneously into a critical nucleus, corresponding to the largest intermediates. As discussed in the main text, we set the $\Delta G_{LLPS}$ zero and regarded the LLPS nuclei as part of the droplet phase. Since oligomers as fibrillization intermediates are structurally distinct from ordered fibrils, we treated them separately and assigned a high energy barrier to account for the nucleated conformational changes. Next, we discuss the derivation of all the net proteins flux terms between different states as used in Eqs. 1-6 in the main text. For the modeling of LLPS, here is the breakup of different kinetic processes:

_Droplet nucleation and disintegration_. Following the classical nucleation theory (CNT), the droplet nucleation rate is given by $k_{dn}M^{n_d}$, where $k_{dn}$ is the nucleation rate constant, M is the monomer concentration. The rate constant, $k_{dn}$, is determined by the pre-equilibration approximation(1), $k_{dn}= k_{d+}(k_{d+}/k'_{d-})^{n_d-2}$, where $k_{d+}$ is droplet association rate and $k'_{d-}$ is the droplet intermediate dissociation rate (**Fig. 1C**). When a monomer dissociates from droplet nucleus/intermediates, these smallest droplets disappear and disintegrate into monomers. This disintegration process is characterized by a rate of change of the droplet number concentration, denoted as, $k'_{d-}\{n_d\}$, where $\{n_d\}$ signifies the number concentration of the droplet nucleus.

_Droplet growth and shrinkage_. The droplets grow or shrink through the monomer addition or detachment, governed by addition and dissociation rate constants, $k_{d+}$ and $k_{d-}$ respectively.

Based on the above discussed processes, the rate of change in the number concentration of droplet nucleus is given by

$$\frac{d\{n_d\}}{dt} = k_{dn} M^{n_d} - k_{d+}\{n_d\}M + k_{d-}\{n_d+1\} - k'_{d-}\{n_d\}. \tag{S1}$$

The rate of change in the number concentration of larger droplets is given by

$$\frac{d\{n_d+1\}}{dt} = k_{d+}\{n_d\}M - k_{d-}\{n_d+1\} - k_{d+}\{n_d+1\}M + k_{d-}\{n_d+2\}, \tag{S2}$$

$$\frac{d\{n_d+2\}}{dt} = k_{d+}\{n_d+1\}M - k_{d-}\{n_d+2\} - k_{d+}\{n_d+2\}M + k_{d-}\{n_d+3\}, \tag{S3}$$

$$\frac{d\{n_d+3\}}{dt} = k_{d+}\{n_d+2\}M - k_{d-}\{n_d+3\} - k_{d+}\{n_d+3\}M + k_{d-}\{n_d+4\}, \tag{S4}$$

and so on. Here, $\{n_d + i\}$ denotes the number concentration of droplets of size $n_d$+i with the integer i ranging from 0 to ∞. The rate of change of the total droplet number concentration $C_D$ can be obtained by summing the rate equations above for all sized droplets,

$$\frac{dC_D}{dt} = k_{dn} M^{n_d} - k'_{d-}\{n_d\}. \quad \text{(S5)}$$

Similarly, the rate of change of the total droplet mass concentration can be obtained by multiplying each equation by their respective droplet size and summing them

$$\frac{dD}{dt} = n_d k_{dn} M^{n_d} + (k_{d+} M - k_{d-}) C_D + k_{d-} \{n_d\} - n_d k'_{d-} \{n_d\}. \quad \text{(S6)}$$

The rate of change of monomer concentration related to droplet formation thus equals to:

$$\frac{dM}{dt} = -n_d k_{dn} M^{n_d} - (k_{d+} M - k_{d-}) C_D - k_{d-} \{n_d\} + n_d k'_{d-} \{n_d\}. \quad \text{(S7)}$$

The net flux of proteins from the solution to droplets as noted as $\mathbf{J_{M \to D}}$ in Eqs. 1-6 equals to

$$J_{M \to D} = n_d k_{dn} M^{n_d} + (k_{d+} M - k_{d-}) C_D + k_{d-} \{n_d\} - n_d k'_{d-} \{n_d\}.$$

Next, we examine different kinetic processes of fibrillization. Both oligomer and fibril can form in both heterogeneous protein phases. We treated droplets of different sizes as a totality for oligomerization and fibrillization. However, due to rapid monomer-droplet equilibration, oligomerization and fibrillization in the droplets do not reduce the high protein concentration but rather decrease the total droplet mass or volume. With shrinking total droplet volume, the oligomers and fibrils initially formed in the droplets become partially exposed to the monomer solution, and grew by the addition of proteins in both protein-rich and poor phases. We used a mean-field approach to model this effect and assumed that the droplet-depletion induced exposure is equally applied to all oligomers and fibrils initially formed in droplets. We applied the volume fraction of total droplets $D/D_{max}$ to model the average exposure, where D is droplet mass concentration and $D_{max}$ denotes the maximum protein concentration of droplets reached during the initial monomer-droplet pre-equilibrium. We also assumed that the disintegration of oligomers initially nucleated within droplets led to added protein monomers partitioned into both phases according to the fraction of solvent-exposure.

<u>*Oligomer nucleation and disintegration*</u>. For simplicity, we treated droplets of different sizes as a single phase for fibrillization. Following CNT, the rate at which the oligomers form in solution phase is $k_{LD,fn} M^{n_f}$, where $k_{LD,fn}$ is the oligomer nucleation rate constant in the solution, $n_f$ is oligomer nucleation size. The rate at which the oligomers form inside the droplets is $k_{HD,fn} \rho^{n_f}$, where $k_{HD,fn}$ is the oligomer nucleation rate constant inside the droplets, and ρ is droplet density. Since droplets only occupy a small fraction of total solution, the rate of loss of droplet mass concentration due to oligomerization inside equals to $k_{HD,fn} \rho^{n_f}(v_D/v)$, where $v_D$ is the droplet volume, v is the total solution volume, and $v_D/v$ is the volume fraction of droplets. Since $\rho v_D/v$ equals to the droplet mass concentration, the reducing rate of droplet mass concentration due to oligomerization becomes $k_{HD,fn} \rho^{n_f - 1} D$. The oligomer nucleation rate constants $k_{LD/HD,fn}$ with the subscript denoting amyloid aggregation in either low-density solution or high-density droplets can be determined by using fast equilibration approximation, $k_{LD/HD,fn} = k_{LD/HD,f+} ( k_{LD/HD,f+}/k'_{LD/HD,f-} )^{n_f - 2}$, where $k_{LD/HD,f+}$ are the fibrillization association rates and $k'_{LD/HD,f-}$ are the fibrilization intermediate or oligomer dissociation rates.

An oligomer in the solution phase disintegrates into monomers within the solution. The disintegration rate of oligomer originally nucleated in the solution phase is given by $k'_{LD,f-}\ \{n_f\}_{LD}$, where $\{n_f\}_{LD}$ is oligomer concentration in the solution. However, oligomers nucleated inside the droplet may disintegrate and release monomers into both phases. The fraction of the oligomer buried inside the droplet is approximated by $\{n_f\}_{HD}\ \frac{D}{D_{max}}$, while the remaining fraction, $\{n_f\}_{HD}\ (1-\frac{D}{D_{max}})$, is exposed to the solution, where $\{n_f\}_{HD}$ is oligomer concentration nucleated within the droplets. Therefore, the rate at which oligomers, nucleated originally inside the droplet, disintegrate into monomers within the droplets is $k'_{HD,f-}\ \{n_f\}_{HD}\ \frac{D}{D_{max}}$, while the rate at which oligomers, nucleated originally inside the droplet, disintegrated into monomer in the solution is $k'_{LD,f-}\ \{n_f\}_{HD}\ (1-\frac{D}{D_{max}})$.

Taken together, the net flux of proteins from the droplet phase into oligomers remained buried inside the droplets corresponds to $\mathbf{J_{D\rightarrow O_{HD,buried}}}$ in Eqs. 1-6,

$$J_{D\rightarrow O_{HD,buried}} = n_f\, k_{HD,fn}\, \rho^{n_f-1}\, D - n_f\, k'_{HD,f-}\, \{n_f\}_{HD}\, \frac{D}{D_{max}}.$$

The net flux of proteins from the monomer solution into oligomers formed in the solution, denoted as $\mathbf{J_{M\rightarrow O_{LD}}}$ in Eqs. 1-6, equals to

$$J_{M\rightarrow O_{LD}} = n_f\, k_{LD,fn}\, M^{n_f} - n_f\, k'_{LD,f-}\, \{n_f\}_{LD},$$

and the net flux of proteins from the monomer solution into oligomers that are initially formed in the droplets but now partially solution-exposed, denoted as $\mathbf{J_{M\rightarrow O_{HD,exposed}}}$, equals to

$$J_{M\rightarrow O_{HD,exposed}} = -n_f\, k'_{LD,f-}\, \{n_f\}_{HD}(1 - \frac{D}{D_{max}}).$$

*Fibril formation and growth*. In our model, an oligomer of size $n_f$ grew into a fibril seed of size $n_f + 1$ by monomer addition while undergoing a nucleated conformational conversion. In the solution phase, the formation rate of fibril seeds converted from oligomers is given by $(k_{LD,f+}\ M\ \{n_f\}_{LD} - k_{LD,f-}\ \{n_f+1\}_{LD})e^{-\Delta G_{LD}}$, where $\{n_f\}_{LD}$ is the oligomer concentration in the solution, $\{n_f+1\}_{LD}$ is the concentration of smallest sized fibrils or fibril seeds, $\Delta G_{LD}$ is the oligomer conformational conversion free-energy barrier in the solution. These smallest fibrils of size $n_f + 1$ continue to grow as more monomers being incorporated. Following the same procedure used to calculate droplet growth discussed above, the growth rate of fibrils formed in the solution is given by $(k_{LD,f+}\ M - k_{LD,f-})\ C_{F,LD} + k_{LD,f-}\ \{n_f+1\}_{LD}$ where $C_{F,LD}$ is the total fibril number concentration in the solution phase. The rate of change of $\mathbf{C_{F,LD}}$ equals to

$$\frac{dC_{F,LD}}{dt} = (k_{LD,f+}\ M\ \{n_f\}_{LD} - k_{LD,f-}\ \{n_f+1\}_{LD})e^{-\Delta G_{LD}}. \qquad \text{(S8)}$$

Considering the partial solvent-exposure of oligomers initially formed in the droplet phase, the formation rate of fibril seeds converted from oligomers by the association of droplet proteins in the buried portion is given by $(k_{HD,f+}\ \rho\ \{n_f\}_{HD} - k_{HD,f-}\ \{n_f+1\}_{HD})\ e^{-\Delta G_{HD}}\ \frac{D}{D_{max}}$, and the formation rate of fibril seeds from oligomers by monomer addition in the solution in the exposure portion is $(k_{LD,f+}\ M\ \{n_f\}_{HD} - k_{LD,f-}\{n_f+1\}_{HD})\ e^{-\Delta G_{LD}}\left(1-\frac{D}{D_{max}}\right)$, where $\{n_f\}_{HD}$ is the oligomer concentration in the droplets, $\{n_f+1\}_{HD}$ is the concentration of smallest sized fibrils, $\Delta G_{HD}$ and

$\Delta G_{LD}$ are the oligomer conformational conversion free-energy barriers in the droplet and solution phases, respectively. The net growth rate of fibrils initially formed within the droplets due to the addition of droplet proteins in the buried portion is given by $\left(k_{HD,f+}\,\rho - k_{HD,f-}\right) C_{F,HD} \frac{D}{D_{max}} + k_{HD,f-}\,\{n_f + 1\}_{HD} \frac{D}{D_{max}}$ where $C_{F,HD}$ is the fibril number concentration in the droplet phase. The rate of change of $\mathbf{C_{F,HD}}$ equals to

$$\frac{dC_{F,HD}}{dt} = (k_{HD,f+}\,\rho\,\{n_f\}_{HD} - k_{HD,f-}\,\{n_f + 1\}_{HD})\, e^{-\Delta G_{HD}} \frac{D}{D_{max}} + (k_{LD,f+} M\,\{n_f\}_{HD} - k_{LD,f-}\{n_f + 1\}_{HD})\; e^{-\Delta G_{LD}} \left(1 - \frac{D}{D_{max}}\right). \tag{S9}$$

The net growth rate of these fibrils due to the addition of solution proteins in the solution exposed portion is given by $\left(k_{LD,f+}\, M - k_{LD,f-}\right) C_{F,HD} \left(1 - \frac{D}{D_{max}}\right) + k_{LD,f-}\,\{n_f + 1\}_{HD} \left(1 - \frac{D}{D_{max}}\right)$.

Together, the net flux of proteins from oligomer to fibril conversion in the solution phase represented by $\mathbf{J_{O_{LD} \to F_{LD}}}$, equals to

$$J_{O_{LD} \to F_{LD}} = n_f\,(k_{LD,f+}\, M\,\{n_f\}_{LD} - k_{LD,f-}\,\{n_f + 1\}_{LD})\, e^{-\Delta G_{LD}}.$$

The net flux of proteins for fibril growth due to monomer addition in the solution phase as represented by $\mathbf{J_{M \to F_{LD}}}$ equals to

$$J_{M \to F_{LD}} = (k_{LD,f+}\, M\,\{n_f\}_{LD} - k_{LD,f-}\,\{n_f + 1\}_{LD})\, e^{-\Delta G_{LD}} + (k_{LD,f+}\, M - k_{LD,f-})\, C_{F,LD} + k_{LD,f-}\,\{n_f + 1\}_{LD}.$$

The net flux of proteins during oligomer to fibril conversion within the droplet phase as represented by $\mathbf{J_{O_{HD} \to F_{HD}}}$ equals to

$$J_{O_{HD} \to F_{HD}} = n_f\,(k_{HD,f+}\,\rho\,\{n_f\}_{HD} - k_{HD,f-}\,\{n_f + 1\}_{HD})\, e^{-\Delta G_{HD}} \frac{D}{D_{max}} + n_f\,(k_{LD,f+}\, M\,\{n_f\}_{HD} - k_{LD,f-}\{n_f + 1\}_{HD})\, e^{-\Delta G_{LD}} \left(1 - \frac{D}{D_{max}}\right).$$

The net flux of proteins during fibril growth within the droplet phase as represented by $\mathbf{J_{D \to F_{HD,buried}}}$ equals to

$$J_{D \to F_{HD,buried}} = (k_{HD,f+}\,\rho\,\{n_f\}_{HD} - k_{HD,f-}\,\{n_f + 1\}_{HD})\, e^{-\Delta G_{HD}} \frac{D}{D_{max}} + \left(k_{HD,f+}\,\rho - k_{LD,f-}\right) C_{F,HD} \frac{D}{D_{max}} + k_{HD,f-}\,\{n_f + 1\}_{HD} \frac{D}{D_{max}}.$$

The net flux of proteins for the growth of fibrils originally nucleated within droplets by monomer addition in the solution-exposed portion, represented by $\mathbf{J_{M \to F_{HD,exposed}}}$, is

$$J_{M \to F_{HD,exposed}} = (k_{LD,f+}\, M\,\{n_f\}_{HD} - k_{LD,f-}\{n_f + 1\}_{HD})\, e^{-\Delta G_{LD}} \left(1 - \frac{D}{D_{max}}\right) + (k_{LD,f+}\, M - k_{LD,f-})\, C_{F,HD} \left(1 - \frac{D}{D_{max}}\right) + k_{LD,f-}\,\{n_f + 1\}_{HD} \left(1 - \frac{D}{D_{max}}\right).$$

As all the net flux of proteins terms in Eqs. 1-6 defined above, there are 13 total variables, including **M**, **D**, $\mathbf{\{n_d\}}$, $\mathbf{\{n_d{+}1\}}$, $\mathbf{C_D}$, $\mathbf{\{n_f\}_{LD}}$, $\mathbf{\{n_f\}_{HD}}$, $\mathbf{C_{F,HD}}$, $\mathbf{\{n_f{+}1\}_{LD}}$, $\mathbf{C_{F,LD}}$, $\mathbf{\{n_f{+}1\}_{HD}}$, $\mathbf{F_{LD}}$, and $\mathbf{F_{HD}}$. With **Eqs.** S1, S5, S8, and S9 defining the rate of change of $\mathbf{\{n_d\}}$, $\mathbf{C_D}$, $\mathbf{C_{F,HD}}$, and $\mathbf{C_{F,LD}}$ correspondingly, there are a total of 10 equations including **Eqs**. 1-6. In order to solve the differential equations, approximations related to the smallest droplets $\mathbf{\{n_d{+}1\}}$ and smallest fibrils of $\mathbf{\{n_f{+}1\}_{LD}}$, and $\mathbf{\{n_f{+}1\}_{HD}}$ are needed. For instance, in prior models adopting the classical nucleation theories, these smallest aggregates were either assumed not disassociating back to the nucleus (e.g., Powers and Powers (2)) or had the same concentrations as the nucleus (e.g., Serrano et al. (1)). During our analyses, we found that these approximates were valid insofar as the monomer concentration was above the critical concentration of the aggregate. However, when the monomer concentration was below the critical concentration of the aggregate (e.g., for the cases in modeling droplet population below or near the binodal point, or in modeling of fibrillization dissolution under high salt concentrations in **Fig. 4**), these previous approximations failed to capture the equilibrium population of the aggregates. For instance, this can be easily explained by using the example of droplet formation with **Eqs. S2-4**. At the equilibrium or steady state with all the rates of changes equal to zero, the ratio of number concentration of neighboring sized droplets $\{j+1\}/\{j\}$ equals to

$$\{j+1\}/\{j\} = M\,k_{d+}/k_{d-} = M/\Phi_{BN},$$

where $\{j\}$ denotes the number concentration of droplets containing j protein, and j ranges from $n_d$ to $\infty$. Therefore, alternative approximations were needed.

We note that at the monomer-droplet equilibrium, $M/\Phi_{BN}$ should be less than 1 and gradually approach 1 when the total protein concentration increases. As a result, the number concentration of large droplets decreased rapidly with increasing sizes. Therefore, we decided to include additional terms for droplets larger than the nucleus and applied approximation for the largest droplet included. Specifically, we included additional $\mathbf{m_d}$ terms, $\{n_d + i\}$, with i from 1 to $\mathbf{m_d}$, with rate of change as defined by **Eqs. S2-4**. In this study, we set $\mathbf{m_d} = 9$. For the largest droplet, the rate of change equation depends on $\{n_d + \mathbf{m_d}{+}1\}$, and we used the following approximation,

$$\{n_d + m_d + 1\} = \begin{cases} \{n_d + m_d\}, & \text{if } M \le \Phi_{BN} \\ \{n_d + m_d\}\dfrac{M}{\Phi_{BN}}, & \text{if } M < \Phi_{BN} \end{cases}.$$

Similarly, we applied the same approximation for fibrils formed in both phases. We also set the additional terms $\mathbf{m_f} = 9$. For the largest fibrils included in the model, we used the following approximation,

$$\{n_f + m_f + 1\}_{LD} = \begin{cases} \{n_f + m_f\}_{LD}, & \text{if } M \le \Phi_{CR} \\ \{n_f + m_f\}_{LD}\dfrac{M}{\Phi_{CR}}, & \text{if } M < \Phi_{CR} \end{cases},$$

And

$$\{n_f + m_f + 1\}_{HD} = \begin{cases} \{n_f + m_f\}_{LD}, & \text{if } \Phi_{BN} \le \Phi_{CR} \\ \{n_f + m_f\}_{LD}\dfrac{\Phi_{BN}}{\Phi_{CR}}, & \text{if } \Phi_{BN} < \Phi_{CR} \end{cases}.$$

Now with $10+m_d+2*m_f$ total variables and equal number of differential equations, we solve the time-evolution of different species with the initial condition at t = 0 where M = Φ and the concentration of other species zero. In our analysis, we found that the above approximation allowed us to accurately capture the equilibrium partition of proteins in different phases below or above the corresponding critical concentrations.

*Maintaining the monomer-droplet equilibrium during oligomerization and fibrillization.* Compared to monomer-droplet equilibrium, formation of oligomers and fibrils are slow. Since we used the mean-field approximation in treating droplets of all sizes as a totality for oligomerization and fibrillization, the droplet depletion is only reflected on the mass concentration of total droplets, but not the number concentration of $C_D$ and $\{n_d\}$. This may lead to instability of the rate-mass differential equations when the droplet mass is depleted near zero. To mitigate this problem, we proposed to correct the rate of changes of $C_D$ and $\{n_d\}$ to reflect the protein depletion of both monomers and droplets due to oligomerization and fibrillization in two phases, by using the rapid monomer-droplet equilibrium assumption. As discussed above, at equilibrium, $\{j+1\}/\{j\} = M/\Phi_{BN}$ and the number concentration $C_D$ and the mass concentration D of total droplets can be derived by summing all the terms from $j = n_d$ to ∞. Let $M/\Phi_{BN} = \alpha$, we can obtain

$$C_D = \sum_{j=n_d}^{\infty}\{j\} = \{n_d\}/(1-\alpha),$$

and

$$D = \sum_{j=n_d}^{\infty} j\,\{j\} = \{n_d\}(n_d + \alpha - n_d\alpha)/(1-\alpha)^2.$$

Taking derivative of above two equations with respect to time and expressing $dC_D/dt$, $dD/dt$ as the function of $dM/dt$ and $dD/dt$,

$$\frac{dC_D}{dt} = \frac{C_D}{D}\frac{dD}{dt} - \frac{\alpha}{(1-\alpha)(n_f+\alpha-n_f\alpha)}\frac{C_D}{M}\frac{dM}{dt}, \qquad \text{(S10)}$$

and

$$\frac{d\{n_d\}}{dt} = \frac{\{n_d\}}{D}\frac{dD}{dt} - \frac{\alpha(1+n_f+\alpha-n_f\alpha)}{(1-\alpha)(n_f+\alpha-n_f\alpha)}\frac{\{n_d\}}{M}\frac{dM}{dt}. \qquad \text{(S11)}$$

Therefore, after the monomer-droplet equilibrium was achieved and when oligomerization and fibrillization started, we replaced **Eqs. S1** and **S5** with the above two equations and plugged in the rate of changes in M and D due to oligomerization and fibrillization in the corresponding phases as defined in **Eqs. 1-2**. Indeed, the instability of the rate-mass differential equations was avoided using the above approximation.

*Fragmentation.* As discussed in the main text, fragmentation occurs less frequently than monomeric fibril dissociation at the fibril ends. To account this difference, the fragmentation rate, $k_{fragment}$, was introduced with respect to the monomeric fibril dissociation rate $k_{LD/HD,f-}$ by the expression, $k_{fragment} = k_{LD/HD,f-}\exp(-\beta\Delta G_f)$, where $\Delta G_f$ is the free energy difference between fibril fragmentation and monomeric fibril dissociation. Following the same mean-field approach as used in Serrano et al.(1) and Knowles et al.(3), the rate of increase in fibril number concentration due to fragmentation was expressed as $k_{fragment}$ $F_{HD}$ and $k_{fragment}$ $F_{LD}$ for solution phase and droplet phase respectively, by assuming the same fragmentation rates in both phases. In addition, for a longer fibril breaking into two shorter fibrils, the fibril length should be at least twice of the smallest fibril seeds. To account for this effect, we proposed a rescaling function Θ for fragmentation that depended on the mean fibril size, $l = F_{HD/LD}/C_{F,HD/LD}$, with the subscript LD and HD denoting the original phase,

$$\Theta(l) = \begin{cases} 1, & l > 2 * (n_f + 1) \\ \dfrac{l}{n_f + 1} - 1, & 1 < l \leq 2 * (n_f + 1) \\ 0, & l \leq n_f + 1 \end{cases}$$

Therefore, $\Theta(\frac{F_{LD}}{C_{F,LD}})\ k_{LD,f-}\ \exp(-\beta\Delta G_f)\ F_{LD}$ and $\Theta(\frac{F_{HD}}{C_{F,HD}})\ k_{LD,f-}\ \exp(-\beta\Delta G_f)\ F_{HD}$ were then added to **Eqs. S8** and **S9**, correspondingly to model fragmentation.

<u>Secondary nucleation.</u> Secondary nucleation occurs on the lateral surfaces of pre-formed fibrils, where locally enriched monomers due to surface adsorption form oligomers and grow into new fibrils by incorporating additional monomers. For simplicity, we assume that the secondary nucleation followed similar molecular mechanism as in the primary nucleation with $n_f = n_s$. As in Ref.(3), we also assumed that the oligomers fall on the surface for further frill growth such that only the oligomerization processes in both phases needed to be modified in order to model the secondary nucleation.

As discussed in the main text, we introduced a surface adsorption energy term, $-\Delta G_s$, to account for the enrichment of local protein concentration on the surface of preformed fibrils, $\phi_{surface}$, by rescaling with a coefficient $C_s = \exp(\beta\Delta G_s)$ with respect to the bulk concentration $\phi_{bulk}$ – i.e., $\phi_{surface} = C_s\ \phi_{bulk}$. The ***local*** rate of loss of protein monomers on the fibril surface, thus, became $k_{fn}\ \phi_{surface}^{n_s} = C_s^{n_s}\ k_{fn}\ \phi_{bulk}^{n_s} = \exp(n_s\beta\Delta Gs)\ k_{fn}\ \phi_{bulk}^{n_s}$. Since secondary nucleation only occurs locally on the fibril surface, the ***total*** rate of loss of protein concentration due to such secondary nucleation equals to $k_{fn}\phi_{surface}^{n_s}(\frac{v_{FS}}{V})$, where $v_{FS}$ denotes the effective volume with locally enriched proteins around the fibril surface, and V is the total solution volume. For simplicity, the total volume of fibrils $V_F$ is a reasonable approximation for $V_{FS}$, and $\frac{V_F}{V}$ becomes the volume fraction of fibrils. This volume fraction of fibrils equals to $F/\rho_{fibril}$ with F denoting the fibril mass concentration and $\rho_{fibril}$ as the protein density of the fibril. Using the droplet density to approximate fibril density, the total rate of loss of protein monomers due to secondary nucleation equals to $n_s C_s^{n_s}\ k_{fn}\ \phi_{buld}^{n_s}\frac{F}{\rho}$, and the rate of gain of oligomer due to secondary nucleation equals to $C_s^{n_s}\ k_{fn}\ \phi_{buld}^{n_s}\ F/\rho$.

Therefore, for the fibrils initially formed in the solution phase, the rate of gain of oligomer concentration is $C_s^{n_f}\, k_{LD,fn}\, M^{n_f}\, F_{LD}/\rho$. Similarly, the solution exposed portion of those fibrils initially formed in the droplet phase, has the rate of gain of oligomer $C_s^{n_f}\, k_{LD,fn}\, M^{n_f} \frac{F_{HD,LD}}{\rho}\left(1 - \frac{D}{D_{max}}\right)$. For the remaining portion in the droplets, the rate of oligomerization due to secondary nucleation equals to $C_s^{n_f}\, k_{HD,fn}\, \rho^{n_f}\, \frac{F_{HD}}{\rho}\, \frac{D}{D_{max}} = C_s^{n_f}\, k_{HD,fn}\, \rho^{n_f-1}\, F_{HD}\, \frac{D}{D_{max}}$, compared to rate of the primary nucleation of oligomers in the droplet as discussed above, $k_{HD,fn}\, \rho^{n_f-1} D$.

Different from the solution phase, the protein concentration in the droplet phase is already very high and the adsorption of proteins onto the fibril surface could lead to significant perturbation in the partition of proteins remaining in the droplet phases and adsorbed on the fibril surface. Assuming the effective droplet mass equals to D' after the surface adsorption, the total protein mass on the fibril surface is $\frac{F_{HD}}{\rho}\, \frac{D'}{D_{max}}\, (C_s\, \rho) = \frac{D'}{D_{max}}\, C_s F_{HD}$. With the conservation of total droplet mass, D = D'(1 + $\frac{F_{HD}}{D_{max}}\, C_s$), and thus, D' = D/(1 + $\frac{F_{HD}}{D_{max}}\, C_s$). As a result, the updated primary nucleation becomes $k_{HD,fn}\, \rho^{n_f-1} \frac{D}{1+\frac{F_{HD}}{D_{max}} C_s}$. Similar, the secondary nucleation within the droplet phase equals to $C_s^{n_f}\, k_{HD,fn}\, \rho^{n_f-1}\, \frac{F_{HD}}{D_{max}}\, \frac{D}{1+\frac{F_{HD}}{D_{max}} C_s}$. Together, the rate of oligomerization due to primary and secondary nucleation in the droplet phase equals to

$$k_{HD,fn}\, \rho^{(n_f-1)}\, D\, \frac{1+\exp(n_f \beta \Delta Gs)\frac{F_{HD}}{D_{max}}}{1+\exp(\beta \Delta Gs)\frac{F_{HD}}{D_{max}}}.$$

Together, the net flux of proteins from the droplet phase into oligomers remained buried inside the droplets $\mathbf{J_{D \to O_{HD,buried}}}$ is updated as

$$J_{D \to O_{HD,buried}} = n_f\, k_{HD,fn}\, \rho^{(n_f-1)}\, D\, \frac{1+\exp(n_f \beta \Delta Gs)\frac{F_{HD}}{D_{max}}}{1+\exp(\beta \Delta Gs)\frac{F_{HD}}{D_{max}}} - n_f\, k'_{HD,f-}\, \{n_f\}_{HD}\, \frac{D}{D_{max}},$$

while the net flux of proteins from the monomer solution into oligomers in the solution $\mathbf{J_{M \to O_{LD}}}$ is updated as

$$J_{M \to O_{LD}} = n_f\, k_{LD,fn}\, M^{n_f}\, (1 + \exp(n_f \beta \Delta Gs)\, [\frac{F_{LD}}{\rho} + \frac{F_{HD,LD}}{\rho}\left(1 - \frac{D}{D_{max}}\right)]) - n_f\, k'_{LD,f-}\, \{n_f\}_{LD},$$

to account for the effect of secondary nucleation. The corresponding net flux terms in **Eqs. 1-3&5** is correspondingly updated to model secondary nucleation. According to the mean field random network model (4), the concentration of peptide modules within the droplet phase formed by 2 nm peptide (Aβ-peptide) is estimated to be around 20 mM.

**B. Parameterization for the test protein-solvent system**

In the rate-mass equations as discussed in the main text and derived above, although many parameters are introduced, they are interrelated, resulting in only a few independent free parameters. The parameters related to each other are as follows: 1) In droplet nucleation, the ratio $k'_{d-}$ / $k_{d+}$ introduces spinodal concentration ($\Phi_{SN}$) i.e., $K_{d+} = k'_{d-}/\Phi_{SN}$. 2) Monomer-droplet equilibrium gives binodal value ($\Phi_{BN}$), i.e., $K_{d+} = k_{d-}/\Phi_{BN}$. 3) Monomer-fibril equilibrium gives the critical concentration ($\Phi_{CR}$) in the solution below which there is no fibril growth, i.e., $k_{LD,f+} = k_{LD,f-}/\Phi_{CR}$. 4) In fibril nucleation, $k_{f2+}$ and $k'_{f2-}$ are related to each other by the supercritical concentration($\Phi_{SU}$), i.e., $K_{LD,f+} = k'_{LD,f-}/\Phi_{SU}$. 5) Fibrilization rates between droplet phase and solution phase are related as: $k_{HD,fn}\,\rho^{n_f-1} = \sigma\, k_{LD,fn}\,\Phi_{BN}{}^{n_f-1}$, $k_{HD,f+}\,\rho = \sigma\, k_{LD,f+}\,\Phi_{BN}$, $k_{HD,f-} = \sigma\, k_{LD,f-}$ and $k'_{HD,f-} = \sigma\, k'_{LD,f-}$. 4) We have used fast pre-equilibration approximation for droplet nucleation and fibril nucleation(1). Based on that, droplet nucleation rate ($k_{dn}$) and fibril nucleation rate ($k_{LD,fn}$) are approximated as follows: $k_{dn} = k_{d+}(k_{d+}/k'_{d-})^{n_d-2}$ and $k_{LD,fn} = k_{LD,f+}(k_{LD,f+}/k'_{LD,f-})^{n_f-2}$. Based on these relations, we identified only few independent free parameters, which are discussed next.

The experimental observation of phase diagrams (5–10) for various peptides revealed a minimum threshold concentration necessary for LLPS, typically ranging from a few to hundreds of μM. In our test system, we assigned the binodal concentration ($\Phi_{BN}$) value of around 30 μM. Given the rapid nucleation of droplets, the stability of the droplet intermediate is not significantly higher than that of the droplet, we therefore approximated spinodal concentration ($\Phi_{BN}$) around 45 μM. Studies on LLPS and biomolecular condensates revealed that the condensate size depended on protein concentration, interactions, and solution conditions. Observations (11–13) suggested that the smallest condensates may contain just a dozen to a few dozen peptides. Therefore, we tentatively considered the critical size for droplet nucleation to be approximately 15 peptides. Acknowledging previous studies (1, 2, 14–16) that identified or assumed the size of fibril nucleation to be somewhere between 2-10, we chose an appropriate size of oligomer nucleation as 4. For simplicity, we opted to set the critical size of oligomer from secondary nucleation, $n_s$, equal to that of $n_f$ from primary nucleation, ensuring consistency across secondary nucleation.

The diffusion-controlled reaction rate constant ($k_{d-}$) in polymeric solutions varies based on factors like polymer size, solvent properties, and reaction type. For low molecular weight polymers, it is typically around $10^1$-$10^3$ $\mu M^{-1}s^{-1}$, while for high molecular weight polymers, it ranges from $10^{-2}$-$10^1$ $\mu M^{-1}s^{-1}$ (17–19). Taking an appropriate average, we chose 10 $\mu M^{-1}s^{-1}$ for the test system. Experimentally, fibril growth has been observed to continue until the monomer concentration reaches approximately 0.1 nM or even smaller . This threshold concentration ($\Phi_{CR}$ = 0.1nM) is the consequence of the equilibrium between the fibril and monomer states. In previous studies (15, 24), the monomer-fibril addition rate in the solution phase was estimated to be ~$10^{-3}$ $\mu M^{-1}s^{-1}$. However, a monomer-droplet addition rate is in the order of $10^{-1}$ $\mu M^{-1}s^{-1}$. To ensure ample time for droplet-monomer equilibrium and align with the extended lag time observed in experimental fibril seed nucleation, we assumed a monomer addition rate during fibrilization in the solution approximately five orders of magnitude slower than the monomer-droplet addition rate, specifically $k_{LD,f+} = 10^{-6}$ $\mu M^{-1}s^{-1}$. So, $k_{LD,f-} = k_{LD,f+}\ \Phi_{CR} = 10^{-8}s^{-1}$. To introduce the oligomer nucleation barrier, we have estimated the monomer-oligomer dissociation rate as faster than monomer-fibril dissociation, i.e., $k'_{LD,f-} = 700\ k_{f-,\ LD}$, so that supercritical concentration ($\Phi_{SU} = \frac{k'_{LD,f-}}{k_{LD,f+}}$) is about 70 μM.

The oligomer conformational conversion barrier (ΔG) is system-specific and can be influenced by multiple factors including concentration, temperature, pH, ionic strength, molecular interactions, and post-translational modifications. Here, we arbitrarily selected such barrier in droplet ($\Delta G_{HD}$) and in the solution ($\Delta G_{LD}$) to be 6KT. All the above free independent parameters can be tuned for different polymer system to study their unique aggregation kinetics.

#### C. Half-time ($\tau_{50}$) derivation in different concentration regimes

##### 1) Far below supercritical concentration

Under the condition that the total monomer concentration $\Phi_{Tot}$ is significantly below the supercritical concentration $\Phi_{SU}$, a closed-form expression for the time-evolution of monomer concentration during fibrilization as obtained by Powers and Powers(2) is

$$M(t) = \Phi_{Tot}\,[\mathrm{sech}(\,k_{f+}\,t\,(\frac{(n+1)\Phi_{tot}^{(n+1)}}{2\,\Phi_{SU}^{n-1}})^{\frac{1}{2}})]^{\frac{2}{n+1}},$$

by solving the simplified equations (e.g., assuming the fast equilibrium approximation with $O_n = K_{f+}\Phi_{SU}^{n-2}M^n$, oligomer nucleation rate is significantly slower than the fibril growth rate, and fibril dissociation rate is negligible) of

$$\frac{dM}{dt} = -K_{f+}MC_F, \tag{S12}$$

$$\frac{dC_F}{dt} = -\frac{dO_n}{dt} = K_{f+}MO_n = K_{f+}\frac{M^{n+1}}{\Phi_{SU}^{n-2}}, \tag{S13}$$

with the initial condition of $M(0) = \Phi_{Tot}$ and the fibril number concentrations $C_F(0) = 0$. The fibril growth follows $F(t) = \Phi_{Tot} - M(t)$. To account for the nucleation conformation conversion barrier ΔG associated with transition of oligomers to fibrils as proposed by Serrano et al.(1), Eq. S13 becomes

$$\frac{dC_F}{dt} = -\frac{dO_n}{dt} = K_{f+}e^{-\beta\Delta G}\, MO_n = K_{f+}\frac{M^{n+1}}{\Phi_{SU}^{n-2}}e^{-\beta\Delta G}. \quad \text{(S14)}$$

Solivng equations S12, S14 the same away as Powers and Powers(2), the closed form expression for the time evolution of monomer concentration becomes,

$$M(t) = \Phi_{Tot}\,[\mathrm{sech}(\,k_{f+}t\,(\frac{(n+1)\,e^{-\beta\Delta G}\,\Phi_{tot}^{(n+1)}}{2\,\Phi_{SU}^{n-1}})^{\frac{1}{2}})]^{\frac{2}{n+1}}.$$

At half-time ($t = \tau_{50}$), $M(\tau_{50}) = \Phi_{Tot}/2$, we get,

$$\tau_{50} = C(n)k_{f+}^{-1}\, e^{\beta\Delta G/2}\,\Phi_{SU}^{(n-1)/2}\Phi_{Tot}^{-(n+1)/2}. \quad \text{(S15)}$$

where $C(n) = \mathrm{sech}^{-1}[2^{-(n+1)/2}]$.

**2) High above the supercritical concentration**

Under the asymptotic high concentration regime, the initial monomer-oligomer pre-equilibrium has most of the portion partitioned as oligomers and fibrilization is dominated by conversion of oligomers into fibrils. Assuming all monomers quickly form oligomers, the fast monomer-oligomer pre-equilibration approximation(1) has $O_n = M(M/\Phi_{SU})^{n-1}$ and $M = \Phi_{SU}^{(n-1)/n}{O_n}^{1/n}$. Focusing on the rate of oligomer loss, equation S14 equals to

$$\frac{dO}{dt} = -k_{f+}\,\Phi_{SU}^{(n-1)/n}\,{O_n}^{(n+1)/n}e^{-\beta\Delta G}. \quad \text{(S16)}$$

By integrating this equation, the solution is

$$O_n(t) = (\frac{n}{nO_n(0)^{-1/n} + k_{f+}e^{-\beta\Delta G}\,t\,\Phi_{SU}^{(n-1)/n}})^n. \quad \text{(S17)}$$

Since the majority of the total proteins is oligomer, the time evolution of the fibril mass as, $F(t) = \Phi_{Tot} - n\,O_n(t)$, with $O_n(0) \sim \Phi_{Tot}/n$. At half-time ($\tau = \tau_{50}$), $F(\tau) = \Phi_{Tot}/2$, we obtain

$$\tau_{50} = C(n)\,k_{f+}^{-1}\,e^{\beta\Delta G}\,\Phi_{SU}^{(1-n)/n}\Phi_{Tot}^{-1/n}, \quad \text{(S18)}$$

where $C(n) = n^{(n+1)/n}(2^{-1/n} - 1)$.

### Supplementary Figures

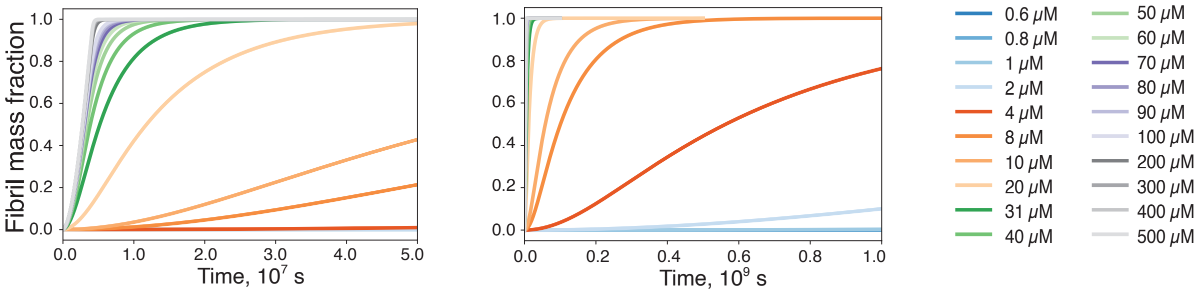


**Figure S1. Time evolution of fibril mass fraction at the twenty different concentrations.** To facilitate efficient curve interpretation, we have adjusted the x-axis range in the two panels. **Fig. 2B** also includes a panel highlighting time evolution of fibril mass fraction at low concentrations (blue lines).

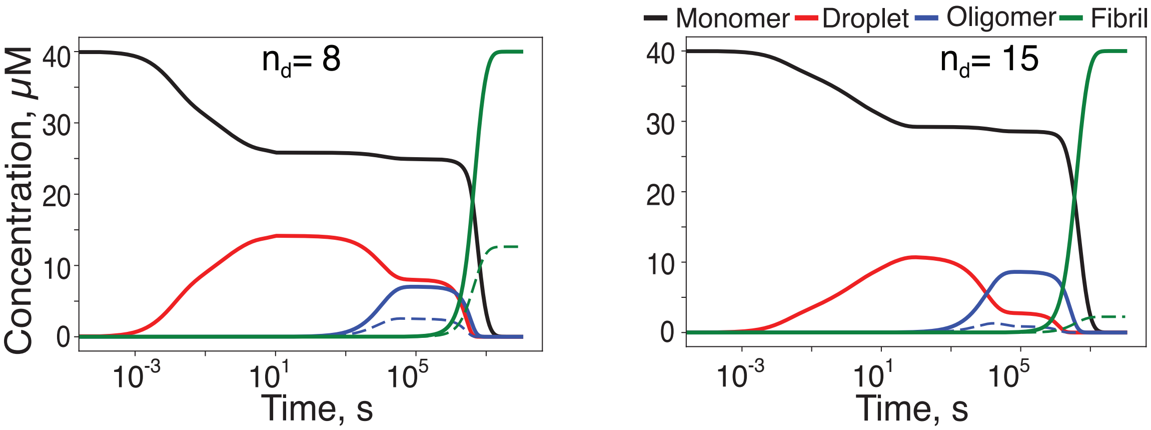


**Figure S2. Time evolution of monomer, droplet, oligomer, fibril state for varying droplet nucleation size ($n_d$) at 40 μM initial protein concentration.**

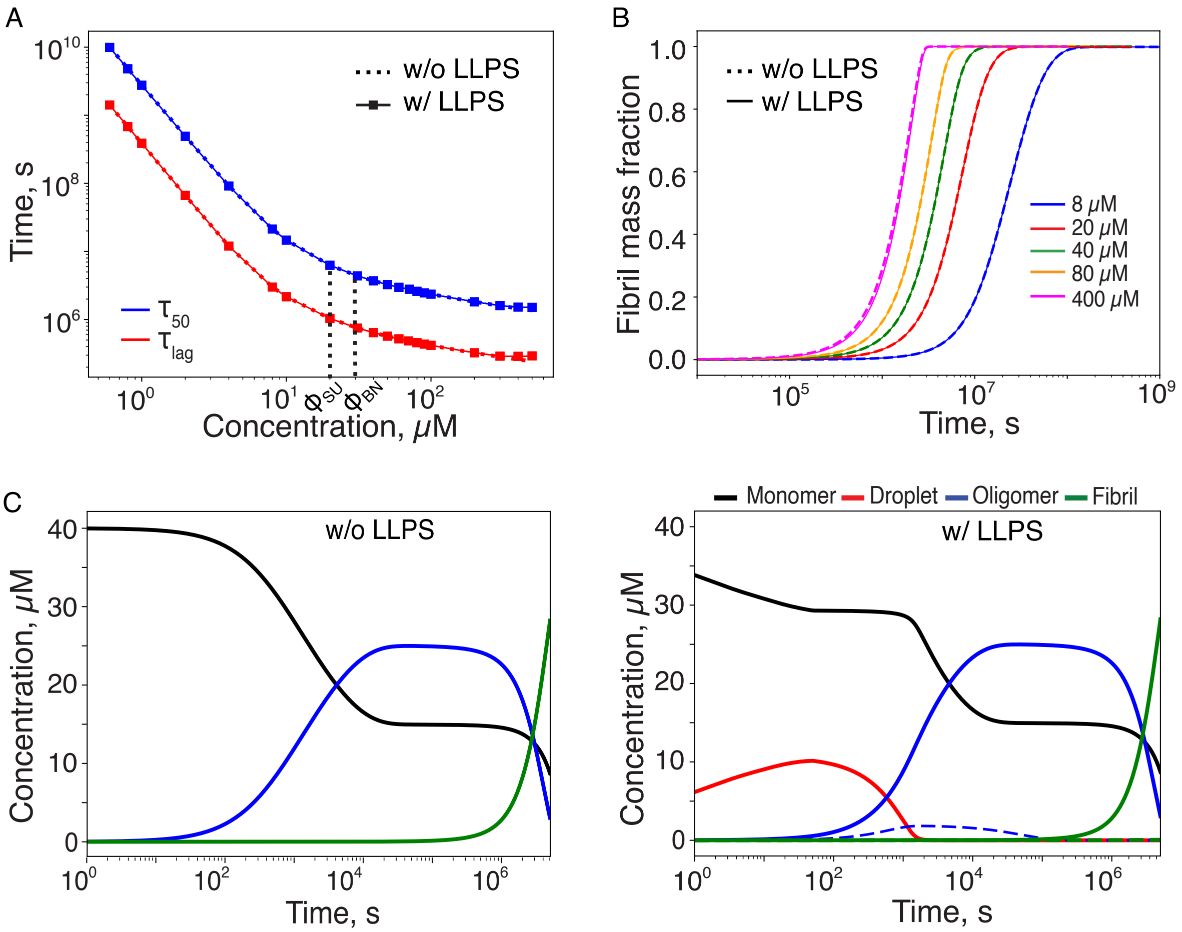


**Figure S3. Impact of oligomer stability on fibril formation kinetics**, when supercritical concentration exceeds binodal point is observed through: **A**) Lag-time ($\tau_{lag}$) and half-time ($\tau_{50}$) for fibril formation as a function of concentration with and without LLPS, **B**) Time evolution of fibril mass fraction at different concentrations with and without LLPS, and **C**) Time evolution of monomers, droplets, oligomers, and fibrils at 40 μM initial protein concentration in both with/without LLPS cases. It is evident that LLPS has no influence on fibril formation kinetics in this situation. The only noticeable effect of LLPS is a slight delay in oligomer nucleation, as monomers first partition into droplets before transitioning into oligomers.

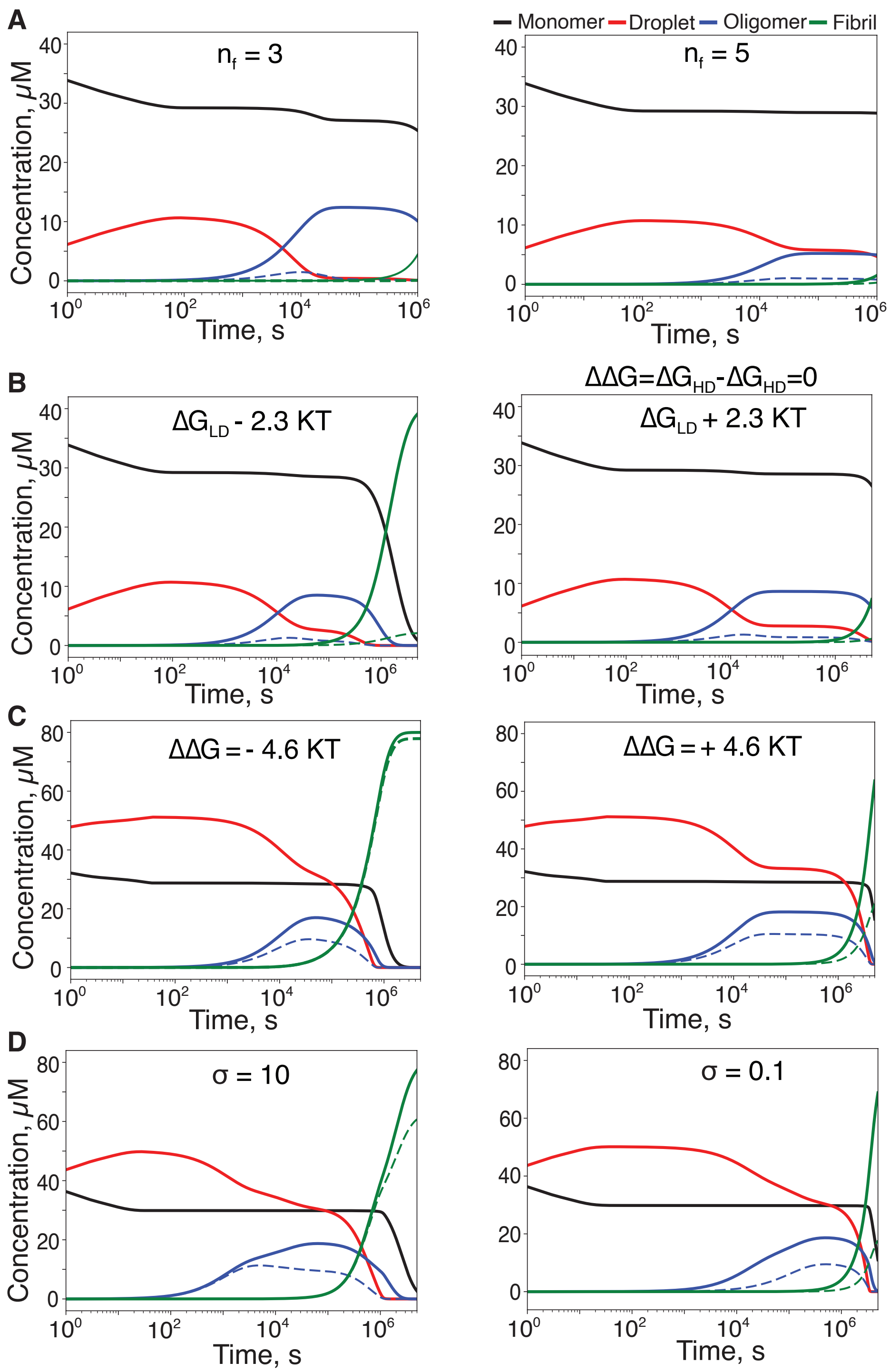


**Figure S4. Time evolution of monomer, droplet, oligomer, fibril state for varying parameter conditions:** A) Fibril nucleation size ($n_f$) at 40 μM initial protein concentration, B) Oligomer conformational conversion barrier equally ($\Delta\Delta G=\Delta G_{HD}-\Delta G_{LD}=0$) in both phases at 40 μM initial protein concentration, C) Difference of oligomer conformational conversion barrier ($\Delta\Delta G$) in two protein phases at 80 μM initial protein concentration, and D) Coefficient σ at 80 μM initial protein concentration.

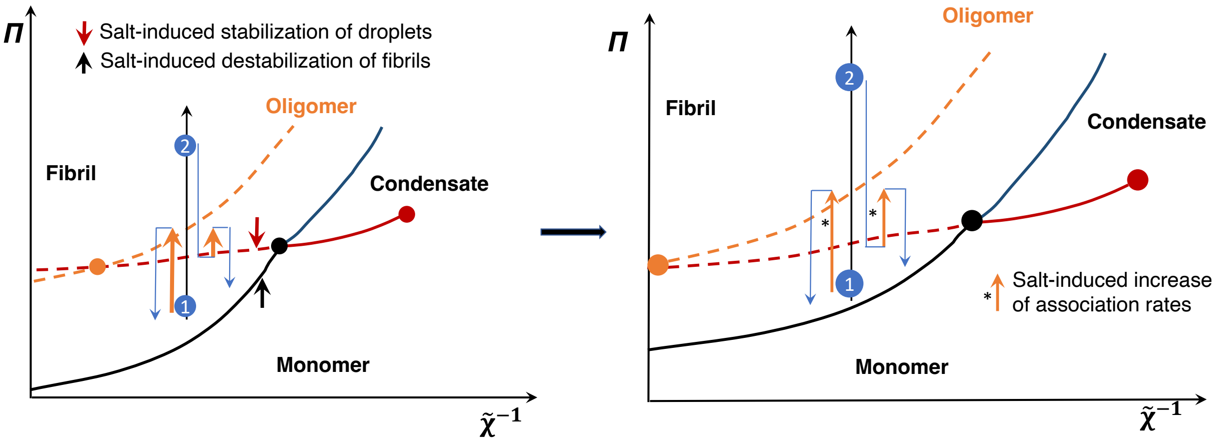


**Figure S5. Phase diagrams of amyloid protein states—monomers, droplets, and fibrils.** The right panel depicts changes in the phase diagram as a result of the presence of salt.

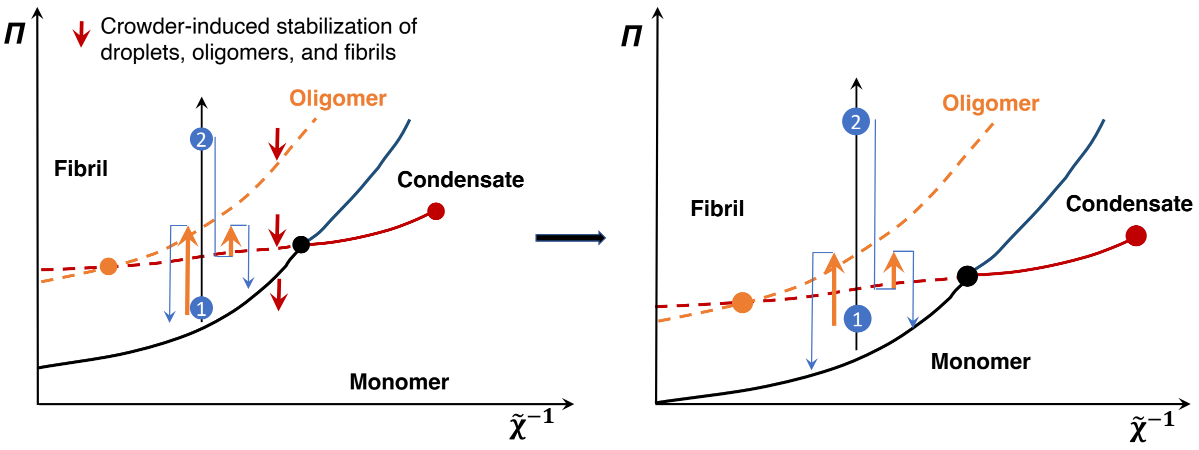


**Figure S6. Phase diagrams of amyloid protein states—monomers, droplets, and fibrils. The right panel depicts changes in the phase diagram as a result of the presence of crodwing agents.**

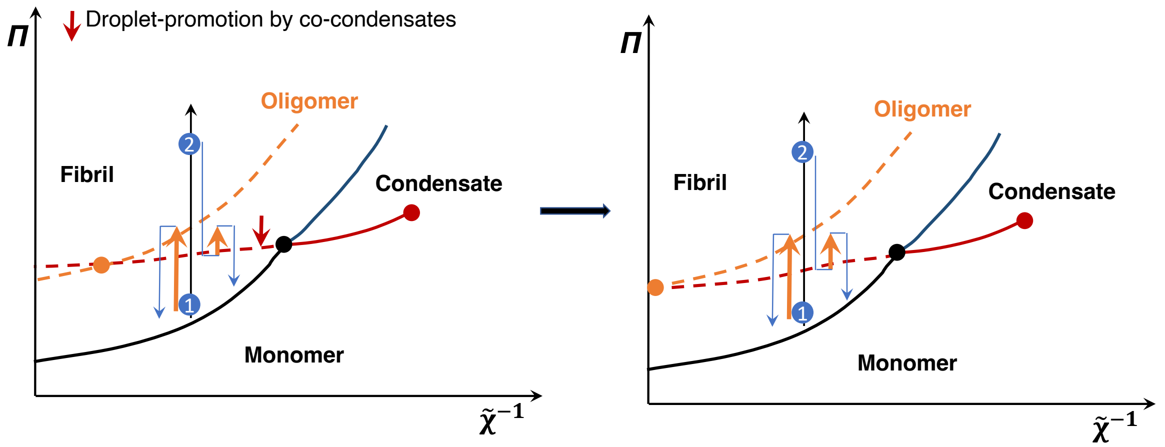


**Figure S7. Phase diagrams of amyloid protein states—monomers, droplets, and fibrils.** The right panel depicts changes in the phase diagram as a result of the presence of co-condensates.

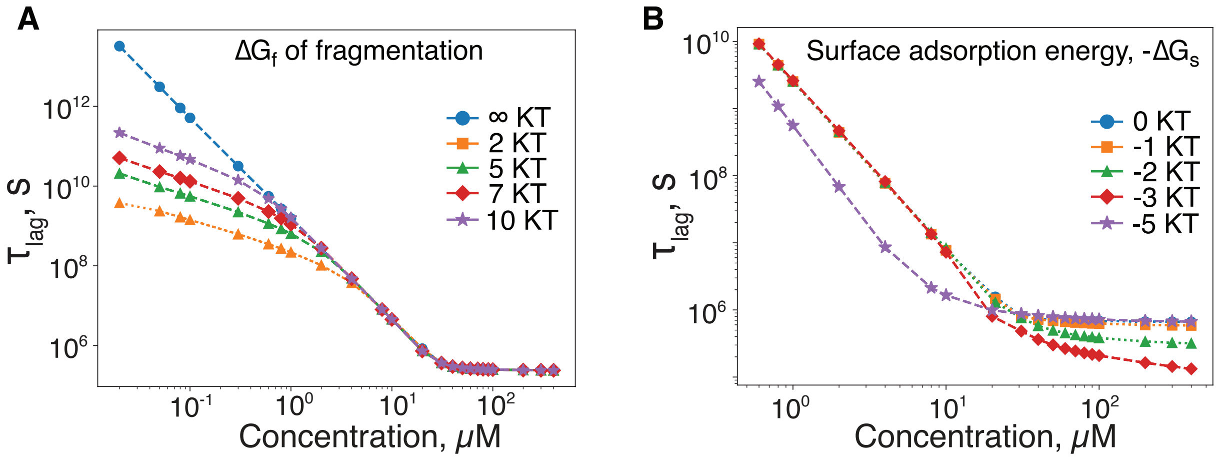


**Figure S8. Effect of fragmentation on fibril formation kinetics.** A) Lag-time ($\tau_{lag}$) for fibril formation as a function of monomer concentration, in comparison with the cases of no fragmentation (blue circles) and at different fragmentation rates. $\Delta G_f$ quantifies how much lower the fragmentation rate is compared to the monomer dissociation rate. B) Effect of secondary nucleation on fibril formation kinetics. Lag-time ($\tau_{lag}$) for fibril formation as a function of monomer concentration, in comparison with the cases of no secondary nucleation (blue circles) or secondary nucleation with different surface adsorption energies ($\Delta G_s$). $\Delta G_s$ captures the propensity of monomer accumulating near the fibril surface to initiate the nucleation.
